# Edinger-Westphal ghrelin receptor signalling regulates binge alcohol consumption in a sex specific manner

**DOI:** 10.1101/2024.03.23.586439

**Authors:** Amy Pearl, Paulo Pinares-Garcia, Arnav Shesham, Xavier Maddern, Roberta G Anversa, Robyn M Brown, Felicia M Reed, William J Giardino, Andrew J Lawrence, Leigh C Walker

## Abstract

**Background:** Rates of risky drinking are continuing to rise, particularly in women, yet sex as a biological variable has been largely ignored. An emerging yet understudied potential component of this circuitry is the central projecting Edinger-Westphal (EWcp), which is made up of two prominent, but distinct cell populations expressing either an array of neuropeptides (including cocaine and amphetamine regulated transcript; CART) or vGlut2 (glutamatergic).

**Methods:** Here, we use a combination of approaches including genetic, molecular biology, behavioural testing, and electrophysiology to understand how the EWcp contributes to alcohol consumption in female versus male mice.

**Results:** Chemogenetic inhibition of EWcp^CART^ cells reduced binge drinking specifically in female, but not male mice. Further, inhibition of EWcp^CART^ cells prevented ghrelin induced drinking, and viral–mediated ghrelin receptor (*Ghsr*) knockdown in the EWcp reduced binge drinking in female, but not male mice. RNAscope revealed *Ghsr* expression across peptidergic (marked by CART) and glutamatergic populations in the EWcp, with neurons from female mice more sensitive to bath application of ghrelin than male mice. Targeted knockdown of *Ghsr* from distinct EWcp populations revealed GHSR signalling on peptidergic, but not glutamatergic cells mediate binge drinking in female mice. Finally, both a GHSR inverse agonist and antagonist delivered directly within the EWcp reduced binge drinking in female mice.

**Conclusions:** These findings suggest the EWcp is a region mediating excessive alcohol bingeing through GHSR actions on peptidergic cells (CART-expressing) in female mice and expand our understanding of the neural mechanism(s) underpinning how the ghrelin system mediates alcohol consumption.

## Introduction

Rates of risky alcohol consumption and alcohol use disorders (AUD) are continuing to rise, particularly in females, where numbers of women engaging in risky drinking and diagnosed with AUD increased over 80% in the last 15 years [1]. The existence of sex differences in alcohol consumption have been widely detailed [2–4]. However, sex as a biological variable has been largely ignored in preclinical research and drug development, with most therapies identified and tested exclusively in male subjects [5]. Thus, prioritising research on the neural mechanisms contributing to AUD in females and understanding sex differences is critical for developing effective treatment strategies.

The presence of binge drinking, defined as a pattern of alcohol consumption that raises blood alcohol levels to 0.08 g/dl (NIAAA) is an early step in the progression of AUD [6]. The putative circuitry mediating this form of excessive alcohol consumption includes the understudied central projecting Edinger-Westphal (EWcp) [7–10]; a structure dense in neuropeptide expression, including cocaine and amphetamine regulated transcript (CART), urocortin (Ucn1) and cholecystokinin (CCK) and sex steroid hormone receptors (estrogen ERα, ERβ and progesterone mPRs) [11, 12]. Recent advancements have shown two major, but discrete populations of neurons within the EWcp exist, and can be broadly defined as peptidergic (expressing CART, Ucn1, CCK) or glutamatergic (vGlut2) [13, 14]. While studies have shown a critical role for this nucleus in alcohol consumption, this has not been well characterised across sexes. Furthermore, while a role for the glutamatergic EWcp in regulating alcohol consumption was recently identified [15], the role of the peptidergic EWcp remains unknown, including the mechanism(s) by which information flowing into the EWcp regulates alcohol-related behaviour more broadly.

Interestingly the EWcp expresses a number of receptors involved in feeding regulation, including dense expression of the ghrelin receptor (growth hormone secretagogue receptor, GHSR) [16, 17]. Indeed, EWcp GHSR expression is striking, and second only to GHSR expression in the well-studied hypothalamic arcuate nucleus (Arc). GHSR is bound by its cognate ligand, ghrelin, a 28-amino acid peptide secreted from the stomach [18]. While this system emerged as an important regulator of energy balance and body weight homeostasis [19] the evidence of the relationship between ghrelin/GHSR and alcohol consumption/seeking is growing (see [20] for review). Preclinical and clinical studies have highlighted a bidirectional relationship between exogenous ghrelin levels and alcohol consumption [20–26], and GHSR antagonists and inverse agonists have shown efficacy in reducing alcohol consumption when administered both peripherally and centrally [24, 27-30]. Further, recent clinical trials have shown safety, tolerability, and some efficacy of GHSR inverse agonist, PF-5190457, to reduce craving in heavy drinking individuals [31, 32].

The central mechanism(s) that underpin ghrelin/GHSRs actions in alcohol consumption and craving are not well understood. Preclinical studies point to a role for GHSR signalling in the laterodorsal tegmental area (LDTg) and ventral tegmental area (VTA) [24, 27, 33]. Interestingly, peripheral administration of the GHSR antagonist, JMV2959, reduces alcohol-induced activation of the EWcp [34], suggesting the EWcp may mediate some of the central actions of GHSRs ability to reduce alcohol consumption.

While most studies examining the ghrelin system have been conducted exclusively in male subjects, sex differences in the effects of ghrelin/GHSR signalling are emerging [35, 36]. Several studies have reported higher circulating levels of ghrelin in females than in males [37–39], and female mice are more sensitive to ghrelin’s effects on feeding [35]. Here we identified sex differences in EWcp CART-expressing cells driving binge-like alcohol consumption. We then tested the hypothesis that GHSR signalling in the EWcp was responsible for these sex specific actions. We found sex differences in *Ghsr* mRNA expression and response to ghrelin in the EWcp, and functionally determined that GHSR signalling on CART-expressing cells in the EWcp drives alcohol consumption through both ligand dependent and independent actions. Together, these data represent a novel mechanism by which ghrelin acts to regulate binge-like alcohol consumption.

## Methods and Materials

For full details please refer to the supplementary information

### Animals

C57BL6J mice (n = 64) were obtained from Animal Resources Centre Australia. Breeding stock of inducible CART-Cre mice (n = 45) [40, 41] and vGlut2-Cre [42] (n = 16, Jackson laboratory stock #016963) were bred in house. iCART-Cre mice were crossed with *Ai14* reporter mice (Jackson laboratory stock #007914) to produce iCART-*Ai14* offspring (n = 14). All studies were performed in accordance with the Prevention of Cruelty to Animals Act (2004), under the guidelines of the National Health and Medical Research Council (NHMRC) and approved by The Florey Animal Ethics Committee.

### Stereotaxic surgery

For viral surgeries 100nL (50 nL/min) of an AAV virus (see Table S1 for virus details) was delivered into the EWcp (M/L + 1.00, A/P −3.40, D/V −4.70 mm on a 15-degree angle) [43]. Cannulae were implanted above the EWcp (M/L + 1.00, A/P −3.40, D/V −4.20 mm on a 15-degree angle) with a dummy projecting 1 mm beyond the cannula tip inserted to maintain patency.

### Drug microinfusion

Vehicle (10% DMSO in 0.9% saline), LEAP2 (5 μg) or JMV2959 (10 μg) were infused (0.25 μL/min, 0.5 uL total) into the EWcp by an automated syringe pump.

### Binge drinking

Mice had voluntary access to 10% (v/v) ethanol, for 2 h, 3 days per week (Monday, Wednesday, Friday) beginning three hours into their dark phase and tested in an extended 4-hour drinking session [4]. For DREADD experiments, CNO or saline were administered 1 h prior to test. To assess ghrelin-induced drinking, access to alcohol was given on the same schedule, but during the light phase when natural ghrelin levels are low [44]. For test sessions, ghrelin was administered 20 minutes prior to test onset, and food removed for the duration of the test.

### qPCR

The EWcp (1.0 mm) and VTA (bilateral 1.0 mm) of each animal were dissected, RNA extracted and RT-qPCR analysis performed as previously described [45, 46]. Primers described in Table S2.

### Immunohistochemistry and Fluorescent In Situ Hybridisation (FISH)

RNAscope Multiplex Fluorescent Reagent Kit V1 was employed following the manufacturer’s protocol and as per our previous publications [46–49]. See Table S2 for product details.

### Microscopy

Images were acquired with a LSM 780 Zeiss confocal microscope and analysis conducted using image J (National Institutes of Health; RRID:SCR_003070). Cells were deemed positive with >5 puncta present [48, 49].

### Electrophysiology

Baseline data were captured during aCSF bath application, followed by continuous perfusion of aCSF containing Ghrelin (1 µM). Current- and voltage-clamp recordings were performed using an Axon Multiclamp 700B amplifier, Digidata 1440 digitizer, and pCLAMP v10 software. All slice electrophysiology data were analysed using Clampfit v10.7.

### Statistical Analysis

All statistical tests details and results are available in Table S3. Data were analysed using GraphPad Prism (v10) software. Data are expressed as mean ± SEM.

## Results

### 1. EW CART cells mediate binge drinking in a sex specific manner

Given the known role of the EWcp in alcohol consumption, and recent finding that chemogenetic activation of EWcp vGlut2 cells reduces alcohol consumption [15], we sought to determine the role of a distinct population of peptidergic EWcp cells in binge drinking using iCART-Cre mice (Fig 1-C). No differences were observed in weight (*p*>0.05; Fig S1B-C) or training alcohol consumption (*p*>0.05; Fig S1D-E) between control (mCherry) and mice with hM4Di (inhibitory) DREADD injected in the EWcp of either sex. Female mice with mCherry virus showed no difference between CNO and saline administration in either cumulative or total alcohol consumption (*p*>0.05; Fig 1D-E), suggesting no off-target effects of CNO. However, female mice expressing hM4Di DREADDs showed specific reduction in alcohol consumption following CNO administration (*p*<0.01, Fig 1F-G). By contrast, male mice with either mCherry or hM4Di (Fig 1H-I) showed no difference in binge alcohol consumption during test sessions between saline and CNO treatment in both cumulative intake or total consumption (*p*>0.05; Fig 1J-K).

**Figure 1:**
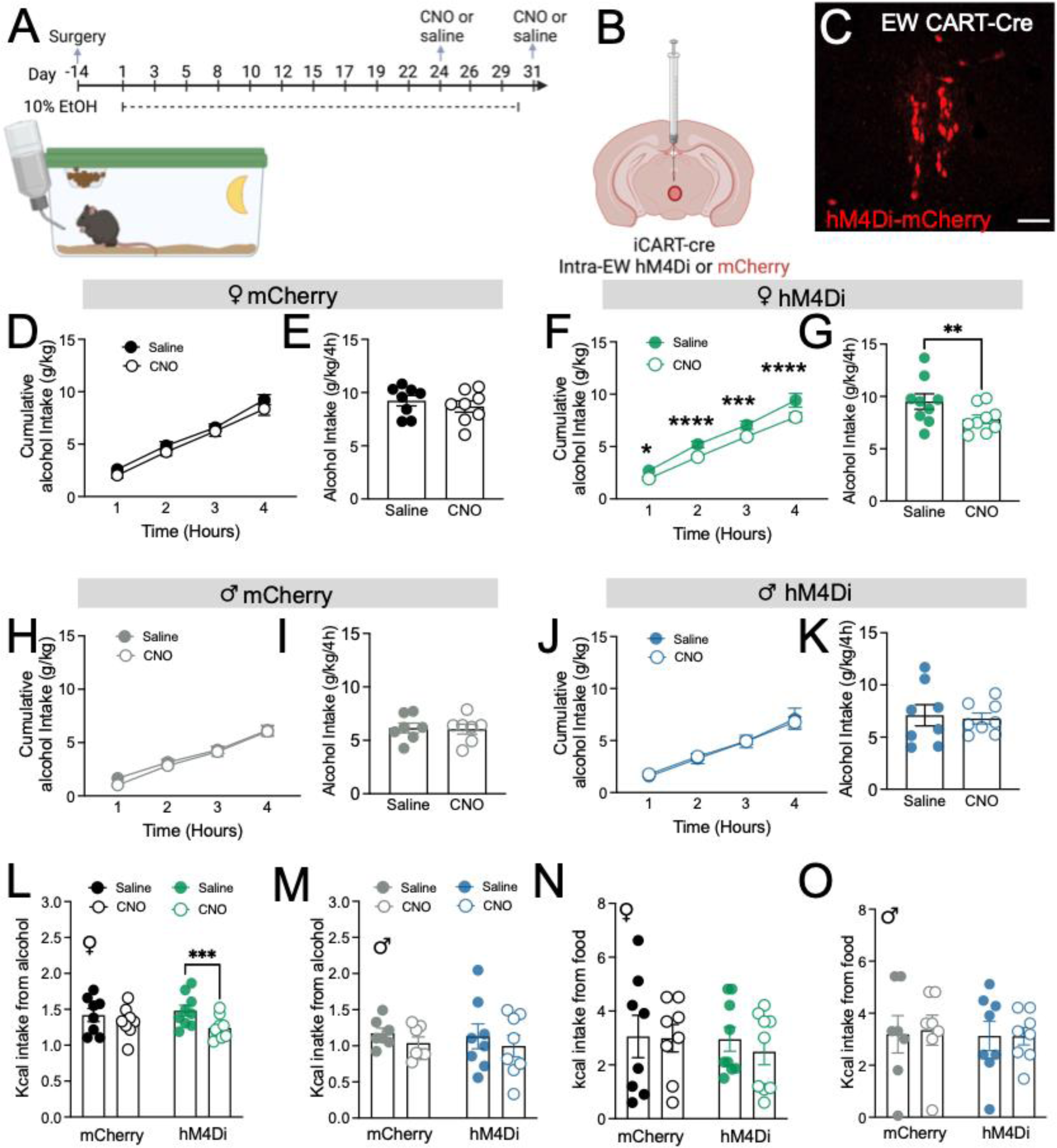
EW^CART^ cells mediate alcohol binge drinking in female, but not male mice. **(A)** Schematic of experimental timeline, **(B)** details of viral approach and **(C)** representative image of viral transduction of EW peptidergic cells in the iCART-Cre mouse. Female mice injected with control virus showed no changes in alcohol consumption following CNO administration both **(D)** in cumulative intake across time or **(E)** total alcohol consumption. However, Female mice injected with hM4Di virus showed a reduction in alcohol consumption following CNO administration both in **(F)** cumulative intake across time and **(G)** total alcohol consumption. Male mice that were injected with control virus showed no changes in alcohol consumption following CNO administration both in **(H)** cumulative intake across time or **(I)** total alcohol consumption. Similarly, male mice injected with hM4Di virus showed no change in **(J)** cumulative or **(K)** total alcohol intake. A significant difference in Kcal intake from alcohol was observed in **(L)** female, but not **(M)** male mice, however no significant difference in Kcal intake from food was observed in **(N)** female or **(O)** male mice. Data expressed as mean ± SEM. n = 8-9 females/group and 7-8 male/group. * *p* < 0.05, ** *p* < 0.01, *** *p* < 0.001, **** *p* < 0.0001. Scale bar = 200 μm.

During the binge drinking test, female hM4Di mice treated with CNO showed a specific reduction in Kcal intake from alcohol (*p*<0.001, Fig L), but not male mice (Fig 1M). Further no difference in Kcal intake from food was observed during the test in either sex (*p*>0.05; Fig 1N-O). Other behaviours were also assessed to determine the specificity of EWcp peptidergic cell inhibition on binge drinking. No difference in water (*p*>0.05; Fig S1F-H) or food intake (*p*>0.05; Fig S1I-J) were observed, nor saccharin preference in either sex (*p*>0.05; Fig S1K-M). Further no difference in anxiety-like behaviours were observed in either sex during the light-dark box procedure (*p*>0.05; Fig S2A-E), no locomotor deficits were observed (*p*>0.05; Fig S2F-G), nor any differences in body temperature (*p*>0.05; Fig S2H-I). Together, these data indicate that EWcp peptidergic neurons specifically reduce alcohol consumption in female mice, without altering other behaviours.

### 2. EW GHSR signalling mediates binge drinking in a sex specific manner

The EW expresses various receptors for feeding-related peptides, including the ghrelin receptor (GHSR), which has been linked to alcohol use [20]. However, the role of these receptors within the EWcp remains unexplored. To assess whether ghrelin signalling in the EWcp regulates alcohol consumption we next trained mice expressing hM4Di DREADD receptors on EW^CART^ cells on a “drinking in the light” procedure, which gave limited access to alcohol for 2 h, 3 times weekly within the light phase when ghrelin levels are low (Fig 2A-C). Ghrelin administration increased alcohol consumption in female, but not male mice (*p*<0.05, Fig 2D-G). In female mice, EW^CART^ DREADD inhibition prevented this escalation in alcohol consumption (*p*>0.05; Fig 2D-G).

**Figure 2.**
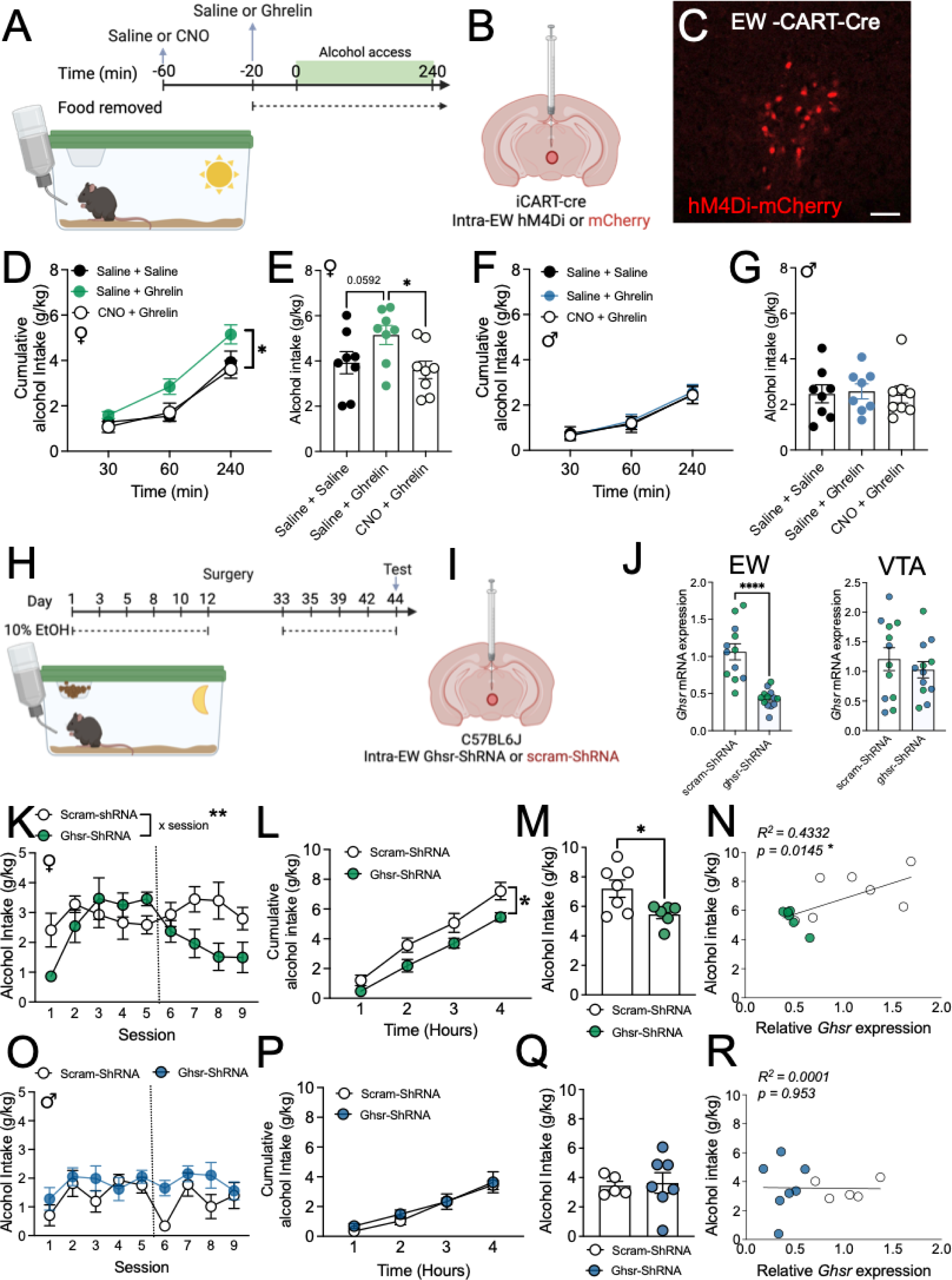
EWcp ghrelin/GHSR signalling regulates alcohol consumption in female, but not male mice. **(A)** Schematic of experimental timeline, **(B)** details of viral approach and **(C)** representative image of viral transduction of EW peptidergic cells in the iCART-Cre mouse. Administration of ghrelin increased **(D)** cumulative and **(E)** total alcohol consumption in female mice, which could be prevented by pre-treatment with CNO. However, ghrelin administration did not alter **(F)** cumulative or **(G)** total alcohol consumption in male mice. **(H)** Schematic of experimental timeline, **(I)** details of viral approach and **(J)** qPCR validation of *Ghsr* mRNA expression in the EW and VTA following shRNA knockdown (males = blue, females = green). **(K)** Training pre- and post-shRNA knockdown showed a significant reduction in alcohol consumption in female mice with shRNA knockdown compared to scram ShRNA following surgery. A significant reduction in alcohol consumption was also observed in *Ghsr*-shRNA treated mice during an extended 4-hour test in both **(L)** cumulative and **(M)** total alcohol intake. **(N)** Alcohol consumption positively correlated with *Ghsr* expression in female mice (open circles = Ghsr-shRNA, closed circles = scram-shRNA. **(O)** Training pre- and post-shRNA knockdown showed no difference in alcohol consumption in male mice with shRNA knockdown compared to scram-ShRNA controls. No difference in alcohol consumption was also observed in male *Ghsr*-shRNA treated mice during an extended 4-hour test in both **(P)** cumulative and **(Q)** total alcohol intake. **(R)** Alcohol consumption did not correlate with *Ghsr* expression in male mice. Data expressed as mean ± SEM. n = 8 females with hM4Di DREADD and 6-7/group for shRNA viruses. n = 8 males with hM4Di DREADDs and 5-7/group for ShRNA viruses. * *p* < 0.05, ** *p* < 0.01, *** *p* < 0.001, **** *p* < 0.0001. Scale bar = 200 μm.

To specifically probe the direct effect of EW^GHSR^ in binge drinking, we disrupted *Ghsr* gene expression in adult mouse brain using an shRNA knockdown approach (Fig 2H-I). RT-qPCR validation showed a specific knockdown of *Ghsr* in the EW, but not the neighbouring VTA in male and female mice (*p*<0.0001, Fig 2J). Female mice injected with Ghsr-ShRNA during training exhibited decreased alcohol consumption compared to mice treated with scram-ShRNA, which showed no change (Fig 2K). This reduction persisted during the test when compared to scram-ShRNA treated mice (*p*<0.05, Fig 2L-M) and EW *Ghsr* expression positively correlated with binge drinking behaviour (*p*<0.05, Fig 2N). No effect of *Ghsr*-ShRNA was observed in male mice (*p*>0.05; Fig 2O-Q), nor did EW *Ghsr* expression correlate with alcohol consumption in males (*p*>0.05; Fig 2R).

### 3. Sex differences in EW Ghsr expression and response to ghrelin

Recent studies have identified two distinct populations of EWcp cells, with distinct neurochemistry (peptidergic vs. glutamatergic) [13, 14]. While dense expression of GHSR has been reported within the EW [16, 17], its distribution across newly defined populations has not been well characterised. Therefore, we conducted RNAscope, RT-qPCR and electrophysiology to explore *Ghsr* expression and function within the EWcp across sexes. Our data show dense expression of *Ghsr* in 99% of peptidergic (*Cartpt+*), and lesser expression in ∼40% of glutamatergic (*Slc17a6+*) cells (Fig 3A-C). Quantification of RNAscope showed a trend towards increased number of *Ghsr*, but not *Cartpt,* or *Slc17a6 (vGlut2)* expressing cells in female mice, compared to male counterparts (Fig 3D). In male mice, 42% of *Ghsr*+ cells were identified as peptidergic, with 49% identified as glutamateric, and the remaining 9% exhibiting neither phenotype. In female mice, 34% of *Ghsr*-positive cells exhibited a peptidergic phenotype, while 61% were identified as glutamatergic, and *Ghsr* expression observed in isolation on 5% of cells (Fig 3E). In line with this, RT-qPCR analysis showed a trend towards greater *Ghsr* expression in female mice (*p*=0.0525, Fig 3G). Further analysis also showed greater expression of the estrogen receptor 1 mRNA (*p*<0.01, ERα, *Esr1,* Fig 3H), and estrogen receptor 2 mRNA (*p*<0.05, ERβ, *Esr2,* Fig 3I), but not membrane bound progesterone receptors (mPRγ, *Parq5;* mPRβ, *Parq8*, *p*>0.05; Fig 3IJ-K) in female mice compared to male counterparts. Whole cell slice recordings revealed no differences in basal electrophysiological properties between sexes (Fig S3). Further, bath application of ghrelin increased firing rate of EWcp peptidergic cells in both female and male mice (*p*<0.001, Fig 3M). However, EW peptidergic cells in female mice were more sensitive to ghrelin, showing a higher firing rate in response to ghrelin compared to male counterparts (*p*<0.05, Fig 3N). A significant decrease in spike amplitude was also observed following bath application of ghrelin in male and female mice (*p*<0.001, Fig 3O), however, there was no difference between sexes (*p*>0.05; Fig 3P). Bath ghrelin application also increased Sag current in female mice significantly more than in male mice (*p*<0.05, Fig S4). Representative traces are shown in Fig 3Q-R for males and females respectively.

**Figure 3.**
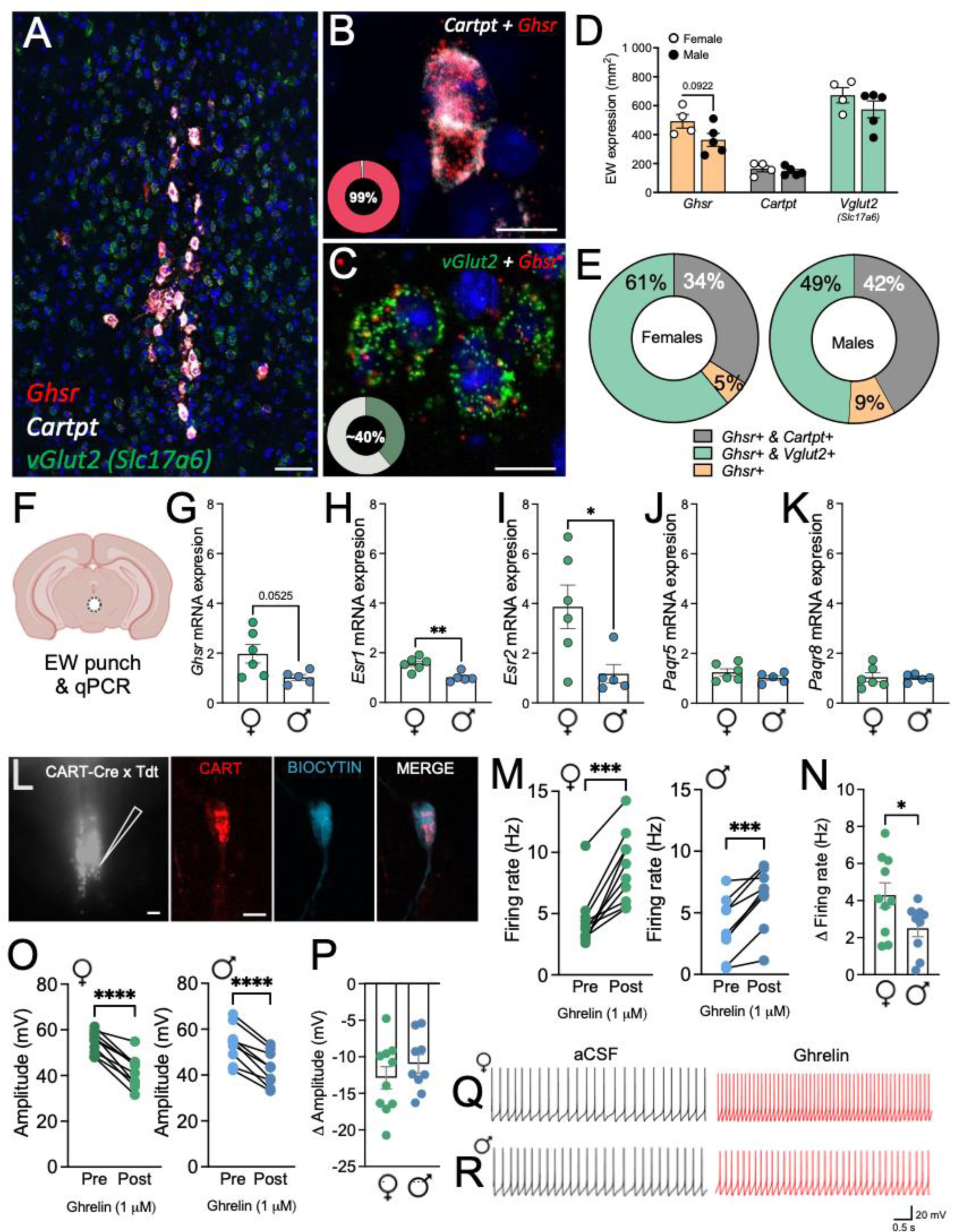
Sex differences are observed in EWcp expression and activity. **(A)** Overview of *Ghsr, Cartpt* and *vGlut2 (Slc17a6)* expression in the Edinger Westphal (EWcp) nucleus. **(B)** High magnification representative microphotograph of *Ghsr* and *Cartpt* expression in the EWcp showed 99% of *Cartpt-*positive cells express *Ghsr.* **(C)** High magnification representative microphotograph of *Ghsr* and *vGlut2* expression in the EW showed 40% of *vGlut2*-positive cells express *Ghsr*. **(D)** Cell density of *Ghsr, Cartpt* and *vGlut2* in male and female mice. **(E)** Donut graph illustrating proportions of *Ghsr-*positive cells that co-express *Cartpt, vGlut2* or neither in male and female mice. **(F)** Schematic showing EW punch location for qPCR analysis between male and female mice. qPCR mRNA analysis for **(G)** the *Ghsr*, estrogen receptors **(H)** *Esr1* (ERα) and **(I)** *Esr2* (ERβ), and progesterone receptors **(J)** *Parq5*, and **(K)** *Parq8*. **(L)** Representative micrograph of endogenous tdTomato expression in the EW of iCART-Tdt mice and confirmation of biocytin filled CART cells. **(M)** Firing frequency at resting membrane potential of EWcp CART+ neurons, showing an increase of the spontaneous firing rate after the application of ghrelin at 1 μM in female (left) and male (right) mice. **(N)** Change in firing rate (delta pre-vs post-ghrelin administration) in male and female mice showed a significant increase in firing frequency in females in response to ghrelin compared to males. **(O)** Amplitude of EWcp CART+ neurons, showing a decrease after the application of ghrelin at 1 μM in female (left) and male (right) mice. **(P)** Change in amplitude (delta pre-vs post-ghrelin administration) in male and female mice showed no significant difference between sexes. Representative traces of EWcp CART+ cells following ghrelin administration in **(Q)** female and **(R)** male mice. Data expressed as mean ± SEM. n = 4-5/sex for RNAscope, 5-6/sex for qPCR and 7-10 cells from 5-7 mice/sex for ephys. * *p* < 0.05, ** *p* < 0.01, *** *p* < 0.001, **** *p* < 0.0001. Scale bar = 200 μm (A & L - overview), 50 μm (B, C & L - overlay).

### 4. Cell-type specific role of GHSR in regulating alcohol consumption

After confirming *Ghsr* is expressed across EW^vGlut2^ and EW^CART^ cells, we next assessed whether *Ghsr* expressed specifically on EW^CART^ or EW^vGlut2^ cells functionally drives alcohol consumption in female mice. To specifically target and disrupt *Ghsr* gene expression in EW subpopulations of the adult mouse brain we used a Cre dependent shRNA approach in iCART-Cre (Fig 4A-C) and vGlut2-Cre (Fig 4G-I) mice. In iCART-Cre mice, *Ghsr*-ShRNA knockdown reduced alcohol consumption during training (Δ alcohol consumption), compared to scram-ShRNA treated mice (*p*<0.05, Fig 4D). This reduction persisted during the test (*p*<0.01, Fig 4E-F). No effect of *Ghsr*-ShRNA was observed in vGlut2-Cre mice compared to scram-ShRNA controls during training (*p*>0.05; Fig 4J), or test (*p*>0.05; Fig 4K-L). We also assessed selective *Ghsr* knockdown on sucrose consumption. *Ghsr*-ShRNA knockdown on EW^CART^ cells did not alter sucrose consumption compared to scram-ShRNA treated mice (*p*>0.05; Fig S5).

**Figure 4.**
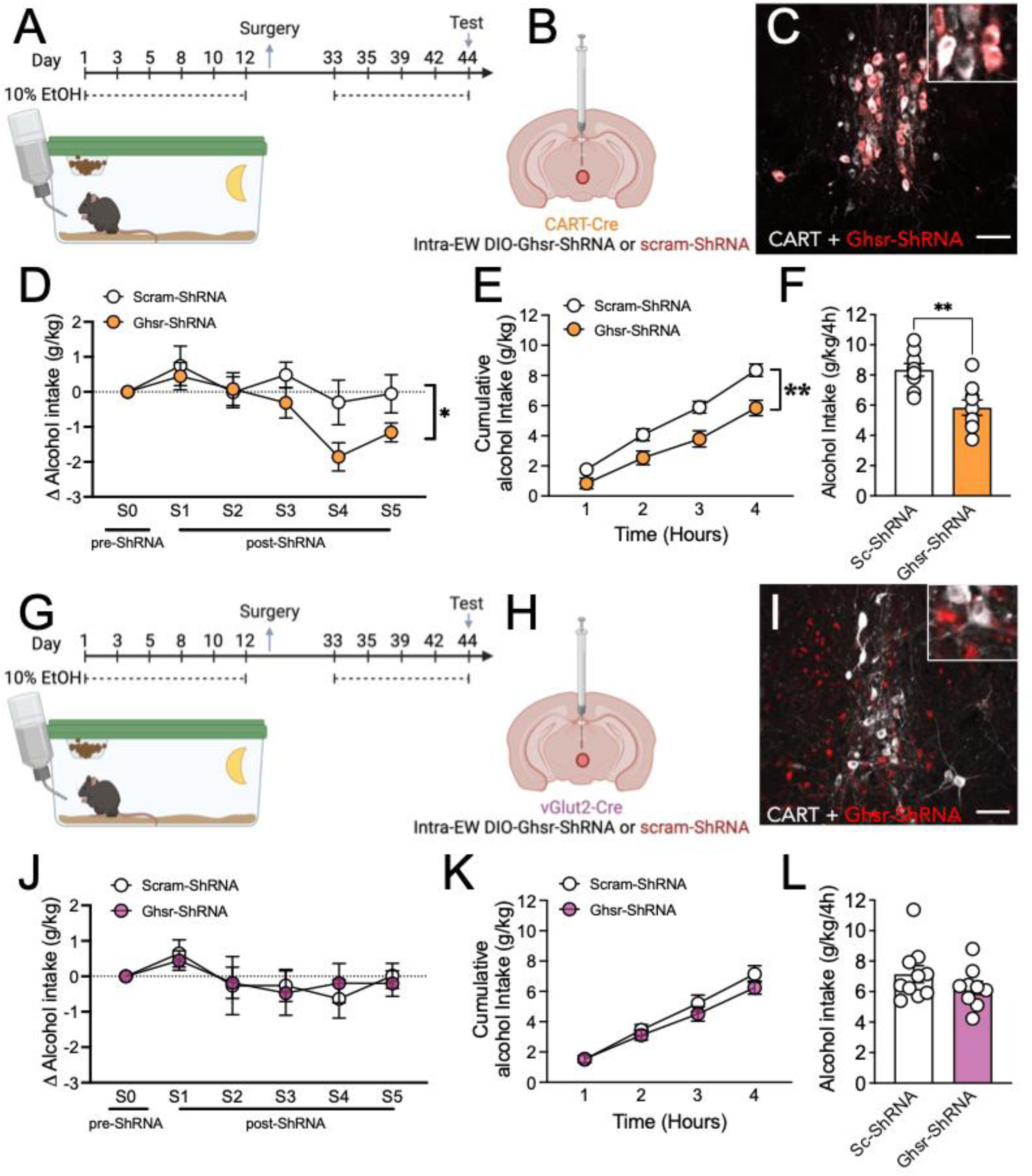
Ghsr acts via EW peptidergic cells to regulate alcohol consumption in female mice. **(A)** Schematic of experimental timeline, **(B)** details of viral approach and **(C)** representative image of viral transduction of EW peptidergic cells in the iCART-Cre mouse. **(D)** GHSR knockdown on EW^CART^ cells reduced alcohol consumption following surgery, **(E)** cumulative alcohol intake and **(F)** total alcohol intake in the 4-hour session. **(G)** Schematic of experimental timeline, **(H)** details of viral approach and **(I)** representative image of viral transduction of EW peptidergic cells in the vGlut2-Cre mouse. **(J)** GHSR knockdown on EW^vGlut2^ cells did not alter alcohol consumption following surgery, **(K)** cumulative alcohol intake or **(L)** total alcohol intake in the 4-hour session. Data expressed as mean ± SEM. n = 8-10/group. * *p* < 0.05, ** *p* < 0.01. Scale bar = 200 μm.

### 5. GHSR1a constitutive and ligand dependent activity drive alcohol consumption in female mice

The *Ghsr* gene encodes two transcripts for GHSR1a and GHSR1b. GHSR1a is a 7-transmembrane GPCR and the cognate receptor for the peptide hormone ghrelin (Sun et al., 2004), whereas the truncated GHSR1b (lacking transmembrane domain 6 and 7) is not bound by ghrelin and is thought to instead regulate trafficking and signalling of GHSR1a within the cell [50]. Interestingly, GHSR1a exhibits unusually high constitutive activity, with ∼50% of its maximal capacity observed in the absence of its agonist, ghrelin [51]. Therefore, we next sought to determine whether the actions of EWcp *Ghsr* knockdown were driven by surface-expressing GHSR1a, and if so whether these actions were via ligand dependent or independent mechanisms. Female mice were microinjected with either vehicle, JMV2959 (a GHSR1a antagonist) or LEAP2 (a GHSR1a inverse agonist) directly within the EW, or adjacent (anatomical control) prior to the binge session (Fig 5A-C). During training, mice with a cannula placed within the EW, or adjacent, showed similar consumption (*p*>0.05; Fig 5D-E) highlighting that cannula implantation *per se* did not alter this behaviour. Administration of JMV2959 or LEAP2 directly within the EW significantly reduced alcohol consumption (Fig 5F). LEAP2 showed significant reduction across all time points (*p*<0.05, Fig 5F), while a significant effect of JMV2959 was noted only after the full 4 h session (*p*<0.05, Fig 5F-G). This was specific to drug administration within the EW, with no effect observed when JMV2959 or LEAP2 were administered adjacent to the EW (Fig 5H-I). Cannula placements are shown in Fig 5J.

**Figure 5.**
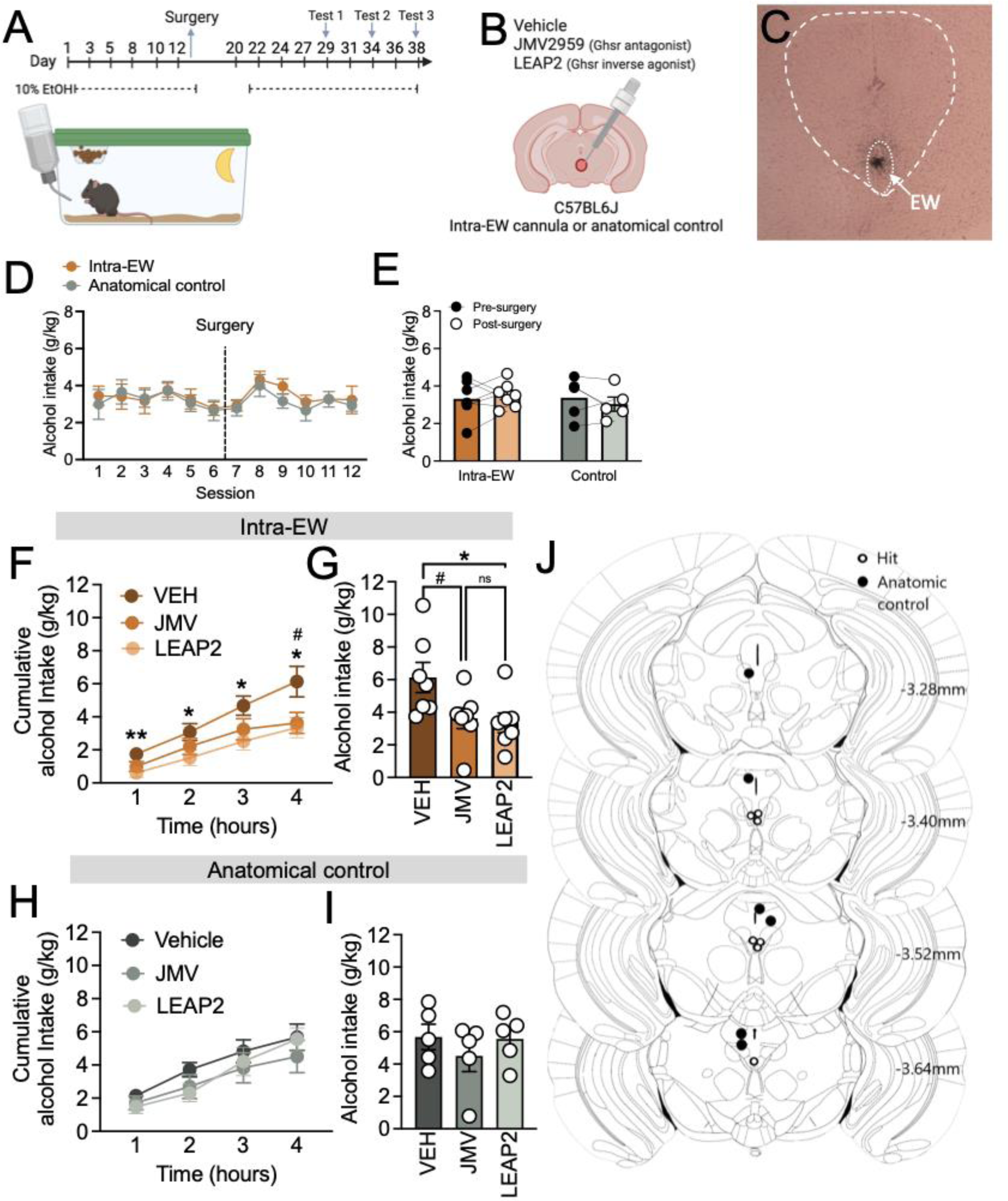
GHSR acts through ligand dependent and independent mechanism to drive alcohol consumption in female mice. **(A)** Schematic of experimental timeline, and **(B)** details of cannulation approach, and **(C)** representative cannula placement in the EW. **(D)** No difference in alcohol consumption was observed across training or **(E)** post-surgery in mice implanted with a cannula in the EW or anatomical controls. **(F)** LEAP2 and JMV2959 administration within the EW reduced cumulative alcohol consumption during the 4-hour test, with LEAP showing significant reduction from hour 1, and JMV2959 after 4 hours. **(G)** LEAP2 and JMV2959 administration within the EW reduced alcohol consumption, however **(H)** administration of LEAP2 and JMV2959 in anatomical controls (adjacent to the EW) did not alter cumulative or **(I)** total alcohol consumption. **(J)** Schematic of cannula placements. Data expressed as mean ± SEM. n = 5-7/group. * *p* < 0.05.

## Discussion

The EWcp is a critical node driving alcohol consumption [7, 10, 52], however the contributions of specific cell populations and mechanisms underpinning this behaviour until now have not been well defined. Our data identify a novel mechanism whereby ghrelin/GHSR1a signalling mediates binge drinking through the peptidergic cells in the EWcp in a sex specific manner.

EWcp peptidergic cells are strongly activated by alcohol [7], in line with this we showed that chemogenetic inhibition of peptidergic EWcp cells reduces alcohol consumption. Interestingly, while alcohol activates the EWcp, specific comparisons of activation between sexes have not been assessed. However, the EWcp has recently been implicated as a connector hub linking regions activated by excessive alcohol consumption in female, but not male mice [53]. In line with this, when inhibiting the EWcp peptidergic cells, we observed a specific reduction of alcohol consumption in female, but not male mice. Recent studies have highlighted two prominent intermingled but separate populations of cells in the EWcp that either synthesise a number of neuropeptides (peptidergic – expressing CART, CCK, Ucn1), or are glutamatergic (expressing vGlut2) [13, 14]. Previous studies have shown *Ucn1* gene knockdown in the EWcp reduces alcohol consumption in extended access procedures, but this did not differ between sexes [8]. Chemogenetic activation of neighbouring vGlut2 cells reduced alcohol consumption in male mice [15], however female mice were not examined. Our data extend these findings to report that chemogenetic inhibition of peptidergic cells reduces alcohol consumption, however, our findings suggest this is specific to female mice, highlighting both behavioural distinction between these populations and potential sex differences in the role of EWcp peptidergic cells in driving alcohol consumption.

The EWcp expresses a number of sex steroid hormone receptors including those for estrogen (ERα & ERβ; Derks et al., 2007) and progesterone (mPRγ & mPRβ; Topilko et al., 2022) which may account for some sex dependent actions driven via the EWcp. Indeed, progesterone reduced firing frequency of EWcp peptidergic cells and ablation of EWcp peptidergic neurons reduced the responsiveness of female mice to progesterone-induced nesting behaviours [13]. We did not observe differences in progesterone receptor mRNA (mPRγ & mPRβ) in the EWcp between sexes in virgin mice, although significant differences in estrogen receptor mRNA expression (ERα & ERβ) were observed. However, the specific roles of sex steroid hormone receptors in the EWcp likely fluctuate based on changes in circulating steroid hormone levels across the estrous cycle and require greater elucidation. Of note, sex differences in levels of both Ucn1 and CART have been reported in in the EW of human suicide victims [54], and studies in rodents show baseline sex differences in Ucn1, CART and nesfatin-1 [55–57] expression in the EW, suggesting the expression of these peptides may be sexually dimorphic in nature.

The EWcp also expresses a number of feeding and arousal related peptide receptors that have gained interest in alcohol research as novel targets for treatment, including orexin/hypocretin [58, 59], leptin [60, 61], and ghrelin[17, 30], however their specific roles within the EWcp have not been characterised. Previous studies have shown increased expression of *Ghsr* mRNA in the EWcp of alcohol preferring male mice (C57BL/6J), compared to a non-preferring strain (DBA/2J) [9]. Our data highlight a sex specific action of *Ghsr* in the EWcp in driving excessive alcohol consumption, whereby ghrelin-induced drinking and viral-mediated knockdown of Ghsr in the EWcp were specific to female mice and driven by interactions with peptidergic cells. Moreover, in female mice, *Ghsr* expression positively correlated with alcohol consumption, highlighting the possibility of a direct relationship between *Ghsr* expression and alcohol intake in females. This is in line with previous reports that show sex differences in circulating levels of ghrelin [37–39], ghrelin levels in abstinent, alcohol dependent individuals [62] and sensitivity to ghrelin induced consummatory behaviour [35].

Little is known about the expression, distribution and function of GHSR in the EWcp of both male and female mice. We identified *Ghsr* mRNA expression on both peptidergic and glutamatergic EWcp populations. Our findings differ from previous reports in *Ghsr*-GFP reporter mice suggesting expression overlapped >90% with the peptidergic marker Ucn1 [17], however this may be due to incomplete reporter expression, or expression levels required to drive recombination and reporter expression in a transgenic mouse line. Indeed, our RNAscope data highlight more dense expression of *Ghsr* mRNA on peptidergic cells, however a large proportion of glutamatergic cells also express *Ghsr* mRNA at lower, but detectable levels. Further, female mice showed a trend towards a greater number of *Ghsr* expressing cells, and *Ghsr* mRNA in the EWcp. Finally, while bath application of ghrelin increased firing rate of peptidergic EWcp cells in both female and male mice, the magnitude of change observed in female mice was significantly higher than male mice, suggesting peptidergic cells in the EWcp of female mice are more sensitive to ghrelin. This is in line with the ability of GHSR to couple to G〈s, and increase intracellular calcium concentrations [18], with bath application of ghrelin increasing the firing rate of cells across the brain including the Arc [63, 64], VTA [65] and central amygdala [66]. However, to our knowledge until now direct comparisons between sexes have not been reported.

Of note, GHSR has high constitutive activity and there is much controversy as to how peripheral ghrelin can access the brain [67, 68]. We observed that both a GHSR1a antagonist (blocking endogenous ghrelin binding) and inverse agonist (inhibiting the GHSR1a directly) when delivered directly into the EWcp reduced excessive alcohol consumption in female mice. These results suggest endogenous ghrelin, not just constitutive activity of the GHSR1a, contributes to alcohol consumption in female mice. Of note, we observed a faster response with the inverse agonist (LEAP2), but the antagonist (JMV2959) had a similar effect when measured over the 4-hour drinking session. The dynamic differences in the time course of effect on alcohol consumption may be explained by differences in direct actions on receptor signalling by the inverse agonist, compared to blockade of ghrelin-induced receptor activity [69, 70] or the specific pharmacokinetics of JMV2959. Recent studies have shown systemic administration of GHSR1a antagonists reduces alcohol consumption in both sexes; however, systemic administration of LEAP2 (inverse agonist) failed to have any effect in either sex [30]. Likely attributed to the undetermined capacity of LEAP2 to penetrate the brain [71], given the same study showed central administration of LEAP2 reduced alcohol consumption [30]. Interestingly, LEAP2 and JMV2959 only reduced consumption in high, but not low drinkers [30]. Given female mice drink alcohol at greater levels than male mice [4], the central actions of GHSR may be more specific to the excessive consumption of alcohol, contributing to the greater effect observed in female mice within this study.

In summary, we highlight a novel neurobiological mechanism that underlies the relationship between ghrelin and alcohol consumption. We identify the EWcp as a novel loci where ghrelin/GHSR1a signalling at peptidergic cells mediates excessive alcohol consumption specifically in female mice. Collectively, our data build upon a growing literature suggesting sex differences in ghrelin/GHSR1a actions in the brain, and elucidate mechanisms underpinning sex differences in excessive alcohol consumption.

## Acknowledgements

We thank the Florey Core Animal Services and Florey Microscopy Facility for their assistance. This project was supported by a Jack Brockhoff Foundation grant and National Health and Medical Research Council (NHMRC) Ideas grant (2002830) awarded to LCW. LCW is also supported by an NHMRC Emerging Leader Fellowship (2008344). AJL is supported by an NHMRC synergy grant (2009851). XJM is supported by an Australian Research Training Program Scholarship. We acknowledge support from the Victorian State Government Operational Infrastructure Scheme.

## Author contributions

AP Data curation; Formal analysis; Investigation; Writing - original draft; Writing - review & editing. PG Data curation; Formal analysis; Investigation; Methodology; Writing - original draft; Writing - review & editing. AS Data curation; Formal analysis; Investigation. XJM Investigation; Writing - review & editing. RGA Investigation; Writing - review & editing. FMR Conceptualization; Writing - review & editing. WJG Conceptualization; Writing - review & editing. AJL Conceptualization; Resources; Supervision; Writing - review & editing. LCW Conceptualization; Data curation; Formal analysis; Funding acquisition; Investigation; Methodology; Project administration; Supervision; Writing – original draft; Writing - review & editing.

## References

1. Grant, B.F., et al., *Prevalence of 12-month alcohol use, high-risk drinking, and DSM-IV alcohol use disorder in the United States*, *2001-2002 to 2012-2013: results from the National Epidemiologic Survey on Alcohol and Related Conditions*. JAMA psychiatry, 2017. 74(9): p. 911–923.

2. Lancaster, F.E. and K. Spiegel, Sex differences in pattern of drinking. Alcohol, 1992. 9(5): p. 415–420.

3. Peltier, M.R., et al., Sex differences in stress-related alcohol use. Neurobiology of stress, 2019. 10: p. 100149.

4. Maddern, X.J., et al., Cocaine and amphetamine regulated transcript (CART) mediates sex differences in binge drinking through central taste circuits. Neuropsychopharmacology, 2024. 49(3): p. 541–550.

5. Lee, S.K., Sex as an important biological variable in biomedical research. BMB reports, 2018. 51(4): p. 167.

6. King, A.C., et al., A prospective 5-year re-examination of alcohol response in heavy drinkers progressing in alcohol use disorder. Biological psychiatry, 2016. 79(6): p. 489–498.

7. Bachtell, R.K., N.O. Tsivkovskaia, and A.E. Ryabinin, Alcohol-induced c-Fos expression in the Edinger-Westphal nucleus: pharmacological and signal transduction mechanisms. Journal of Pharmacology and Experimental Therapeutics, 2002. 302(2): p. 516–524.

8. Giardino, W., et al., Control of chronic excessive alcohol drinking by genetic manipulation of the Edinger–Westphal nucleus urocortin-1 neuropeptide system. Translational psychiatry, 2017. 7(1): p. e1021–e1021.

9. Giardino, W.J., et al., Characterization of genetic differences within the centrally projecting Edinger– Westphal nucleus of C57BL/6J and DBA/2J mice by expression profiling. Frontiers in neuroanatomy, 2012. 6: p. 5.

10. Ryabinin, A.E., et al., High alcohol/sucrose consumption during dark circadian phase in C57BL/6J mice: involvement of hippocampus, lateral septum and urocortin-positive cells of the Edinger-Westphal nucleus. Psychopharmacology, 2003. 165: p. 296–305.

11. Zuniga, A. and A.E. Ryabinin, Involvement of centrally projecting Edinger–Westphal nucleus neuropeptides in actions of addictive drugs. Brain Sciences, 2020. 10(2): p. 67.

12. Pomrenze, M.B., L.C. Walker, and W.J. Giardino, Gray areas: Neuropeptide circuits linking the Edinger-Westphal and Dorsal Raphe nuclei in addiction. Neuropharmacology, 2021. 198: p. 108769.

13. Topilko, T., et al., Edinger-Westphal peptidergic neurons enable maternal preparatory nesting. Neuron, 2022. 110(8): p. 1385–1399. e8.

14. Priest, M.F., et al., Peptidergic and functional delineation of the Edinger-Westphal nucleus. Cell reports, 2023. 42(8).

15. Zuniga, A., et al., Vesicular glutamate transporter 2-containing neurons of the centrally-projecting Edinger-Westphal nucleus regulate alcohol drinking and body temperature. Neuropharmacology, 2021. 200: p. 108795.

16. Zigman, J.M., et al., Expression of ghrelin receptor mRNA in the rat and the mouse brain. Journal of Comparative Neurology, 2006. 494(3): p. 528–548.

17. Spencer, S.J., et al., Ghrelin regulates the hypothalamic-pituitary-adrenal axis and restricts anxiety after acute stress. Biological psychiatry, 2012. 72(6): p. 457–465.

18. Kojima, M., et al., Ghrelin is a growth-hormone-releasing acylated peptide from stomach. Nature, 1999. 402(6762): p. 656-660.

19. Tschöp, M., D.L. Smiley, and M.L. Heiman, Ghrelin induces adiposity in rodents. Nature, 2000. 407(6806): p. 908-913.

20. Farokhnia, M., et al., Ghrelin: From a gut hormone to a potential therapeutic target for alcohol use disorder. Physiology & behavior, 2019. 204: p. 49–57.

21. Calissendorff, J., et al., Inhibitory effect of alcohol on ghrelin secretion in normal man. European journal of endocrinology, 2005. 152(5): p. 743–747.

22. Calissendorff, J., et al., Alcohol ingestion does not affect serum levels of peptide YY but decreases both total and octanoylated ghrelin levels in healthy subjects. Metabolism, 2006. 55(12): p. 1625–1629.

23. Jerlhag, E., Animal studies reveal that the ghrelin pathway regulates alcohol-mediated responses. Frontiers in Psychiatry, 2023. 14: p. 1050973.

24. Jerlhag, E., et al., Requirement of central ghrelin signaling for alcohol reward. Proceedings of the National Academy of Sciences, 2009. 106(27): p. 11318–11323.

25. Leggio, L., et al., Fasting-induced increase in plasma ghrelin is blunted by intravenous alcohol administration: a within-subject placebo-controlled study. Psychoneuroendocrinology, 2013. 38(12): p. 3085–3091.

26. Leggio, L., et al., Intravenous ghrelin administration increases alcohol craving in alcohol-dependent heavy drinkers: a preliminary investigation. Biological psychiatry, 2014. 76(9): p. 734–741.

27. Jerlhag, E., et al., Glutamatergic regulation of ghrelin-induced activation of the mesolimbic dopamine system. Addiction biology, 2011. 16(1): p. 82–91.

28. Gomez, J.L., et al., Differential effects of ghrelin antagonists on alcohol drinking and reinforcement in mouse and rat models of alcohol dependence. Neuropharmacology, 2015. 97: p. 182–193.

29. Gomez, J.L. and A.E. Ryabinin, The Effects of ghrelin antagonists [D-L ys3]-GHRP-6 or JMV 2959 on ethanol, water, and food Intake in C57BL/6J Mice. Alcoholism: Clinical and Experimental Research, 2014. 38(9): p. 2436–2444.

30. Richardson, R.S., et al., Pharmacological GHSR (ghrelin receptor) blockade reduces alcohol binge-like drinking in male and female mice. Neuropharmacology, 2023. 238: p. 109643.

31. Cobbina, E., et al., A population pharmacokinetic analysis of PF-5190457, a novel ghrelin receptor inverse agonist in healthy volunteers and in heavy alcohol drinkers. Clinical pharmacokinetics, 2021. 60: p. 471–484.

32. Lee, M.R., et al., Endocrine effects of the novel ghrelin receptor inverse agonist PF-5190457: Results from a placebo-controlled human laboratory alcohol co-administration study in heavy drinkers. Neuropharmacology, 2020. 170: p. 107788.

33. Edvardsson, C.E., J. Vestlund, and E. Jerlhag, A ghrelin receptor antagonist reduces the ability of ghrelin, alcohol or amphetamine to induce a dopamine release in the ventral tegmental area and in nucleus accumbens shell in rats. European Journal of Pharmacology, 2021. 899: p. 174039.

34. Kaur, S. and A.E. Ryabinin, Ghrelin receptor antagonism decreases alcohol consumption and activation of perioculomotor urocortin-containing neurons. Alcoholism: Clinical and Experimental Research, 2010. 34(9): p. 1525–1534.

35. Yamada, C., et al., Vulnerability to psychological stress-induced anorexia in female mice depends on blockade of ghrelin signal in nucleus tractus solitarius. British journal of pharmacology, 2020. 177(20): p. 4666–4682.

36. Börchers, S., et al., From an empty stomach to anxiolysis: molecular and behavioral assessment of sex differences in the ghrelin axis of rats. Frontiers in Endocrinology, 2022. 13: p. 901669.

37. Johnson, M.L., M.J. Saffrey, and V.J. Taylor, Plasma ghrelin concentrations were altered with oestrous cycle stage and increasing age in reproductively competent Wistar females. Plos one, 2016. 11(11): p. e0166229.

38. Priego, T., et al., Sex-associated differences in the leptin and ghrelin systems related with the induction of hyperphagia under high-fat diet exposure in rats. Hormones and behavior, 2009. 55(1): p. 33–40.

39. Abu-Farha, M., et al., Gender differences in ghrelin association with cardiometabolic risk factors in arab population. International journal of endocrinology, 2014. 2014.

40. Lau, J., et al., CART neurons in the arcuate nucleus and lateral hypothalamic area exert differential controls on energy homeostasis. Molecular metabolism, 2018. 7: p. 102–118.

41. Lau, J., Y.-C. Shi, and H. Herzog, Temperature dependence of the control of energy homeostasis requires CART signaling. Neuropeptides, 2016. 59: p. 97–109.

42. Anversa, R., et al., A paraventricular thalamus to insular cortex glutamatergic projection gates “emotional” stress-induced binge eating in females. Neuropsychopharmacology, 2023. 48(13): p. 1931–1940.

43. Paxinos, G. and K.B. Franklin, Paxinos and Franklin’s the mouse brain in stereotaxic coordinates. 2019: Academic press.

44. Bodosi, B., et al., *Rhythms of ghrelin, leptin, and sleep in rats: effects of the normal diurnal cycle, restricted feeding, and sleep deprivation.* American Journal of Physiology-Regulatory, Integrative and Comparative Physiology, 2004. 287(5): p. R1071–R1079.

45. Walker, L.C., et al., Acetylcholine muscarinic M4 receptors as a therapeutic target for alcohol use disorder: converging evidence from humans and rodents. Biological Psychiatry, 2020. 88(12): p. 898–909.

46. Walker, L.C., et al., Muscarinic M4 and M5 receptors in the ventral subiculum differentially modulate alcohol seeking versus consumption in male alcohol-preferring rats. British journal of pharmacology, 2021. 178(18): p. 3730–3746.

47. Wang, F., et al., RNAscope: a novel in situ RNA analysis platform for formalin-fixed, paraffin-embedded tissues. The Journal of molecular diagnostics, 2012. 14(1): p. 22–29.

48. Viden, A., et al., Organisation of enkephalin inputs and outputs of the central nucleus of the amygdala in mice. Journal of Chemical Neuroanatomy, 2022. 125: p. 102167.

49. Walker, L.C., et al., Cocaine and amphetamine regulated transcript (CART) signalling in the central nucleus of the amygdala modulates stress-induced alcohol seeking. Neuropsychopharmacology, 2021. 46(2): p. 325–333.

50. Mary, S., et al., Heterodimerization with its splice variant blocks the ghrelin receptor 1a in a non-signaling conformation: a study with a purified heterodimer assembled into lipid discs. Journal of Biological Chemistry, 2013. 288(34): p. 24656–24665.

51. Holst, B., et al., High constitutive signaling of the ghrelin receptor—identification of a potent inverse agonist. Molecular endocrinology, 2003. 17(11): p. 2201–2210.

52. Bachtell, R.K., et al., The Edinger-Westphal–Lateral Septum Urocortin Pathway and Its Relationship to Alcohol Consumption. Journal of Neuroscience, 2003. 23(6): p. 2477–2487.

53. Ardinger, C.E., et al., Sex Differences in Neural Networks Recruited by Frontloaded Binge Alcohol Drinking. bioRxiv, 2024: p. 2024.02. 08.579387.

54. Bloem, B., et al., Sex-specific differences in the dynamics of cocaine-and amphetamine-regulated transcript and nesfatin-1 expressions in the midbrain of depressed suicide victims vs. controls. Neuropharmacology, 2012. 62(1): p. 297–303.

55. Derks, N.M., et al., Sex-specific expression of BDNF and CART in the midbrain non-preganglionic Edinger–Westphal nucleus in the rat. Peptides, 2009. 30(12): p. 2268–2274.

56. Derks, N.M., et al., Sex differences in urocortin 1 dynamics in the non-preganglionic Edinger– Westphal nucleus of the rat. Neuroscience research, 2010. 66(1): p. 117–123.

57. Xu, L., et al., Sex-specific effects of fasting on urocortin 1, cocaine-and amphetamine-regulated transcript peptide and nesfatin-1 expression in the rat Edinger–Westphal nucleus. Neuroscience, 2009. 162(4): p. 1141–1149.

58. Emmerzaal, T., et al., Orexinergic innervation of urocortin1 and cocaine and amphetamine regulated transcript neurons in the midbrain centrally projecting Edinger–Westphal nucleus. Journal of Chemical Neuroanatomy, 2013. 54: p. 34–41.

59. Walker, L.C. and A.J. Lawrence, The role of orexins/hypocretins in alcohol use and abuse. Behavioral Neuroscience of Orexin/Hypocretin, 2017: p. 221–246.

60. Bach, P., A. Koopmann, and F. Kiefer, The impact of appetite-regulating neuropeptide leptin on alcohol use, alcohol craving and addictive behavior: a systematic review of preclinical and clinical data. Alcohol and Alcoholism, 2021. 56(2): p. 149–165.

61. Xu, L., et al., Integration of stress and leptin signaling by CART producing neurons in the rodent midbrain centrally projecting Edinger-Westphal nucleus. Frontiers in Neuroanatomy, 2014. 8: p. 8.

62. Wurst, F.M., et al., Gender differences for ghrelin levels in alcohol-dependent patients and differences between alcoholics and healthy controls. Alcoholism: Clinical and Experimental Research, 2007. 31(12): p. 2006–2011.

63. Chen, S.R., et al., Ghrelin receptors mediate ghrelin-induced excitation of agouti-related protein/neuropeptide Y but not pro-opiomelanocortin neurons. Journal of neurochemistry, 2017. 142(4): p. 512–520.

64. Hashiguchi, H., et al., Direct versus indirect actions of ghrelin on hypothalamic NPY neurons. PloS one, 2017. 12(9): p. e0184261.

65. Navarro, G., et al., Complexes of ghrelin GHS-R1a, GHS-R1b, and dopamine D1 receptors localized in the ventral tegmental area as main mediators of the dopaminergic effects of ghrelin. Journal of Neuroscience, 2022. 42(6): p. 940–953.

66. Cruz, M.T., et al., Ghrelin increases GABAergic transmission and interacts with ethanol actions in the rat central nucleus of the amygdala. Neuropsychopharmacology, 2013. 38(2): p. 364–375.

67. Banks, W.A., et al., Extent and direction of ghrelin transport across the blood-brain barrier is determined by its unique primary structure. Journal of Pharmacology and Experimental Therapeutics, 2002. 302(2): p. 822–827.

68. Rhea, E.M., et al., Ghrelin transport across the blood–brain barrier can occur independently of the growth hormone secretagogue receptor. Molecular metabolism, 2018. 18: p. 88–96.

69. Moulin, A., et al., *The 1*, *2*, *4-triazole as a scaffold for the design of ghrelin receptor ligands: development of JMV 2959, a potent antagonist*. Amino acids, 2013. 44: p. 301–314.

70. Wang, J.H., et al., Identifying the binding mechanism of LEAP 2 to receptor GHSR 1a. The FEBS Journal, 2019. 286(7): p. 1332–1345.

71. Lu, X., et al., LEAP-2: an emerging endogenous ghrelin receptor antagonist in the pathophysiology of obesity. Frontiers in Endocrinology, 2021. 12: p. 717544.

