## Supplemental Methods for "Edinger-Westphal ghrelin receptor signalling regulates binge alcohol consumption in a sex specific manner"

Pearl et al.,

**SUPPLEMENTAL METHODS**

*Animals*

C57BL6J mice (n = 64) were obtained from ARC (Animal Resources Australia, WA, Australia) at ~6 weeks of age. Breeding stock of inducible CART-Cre knock in mice (iCART-Cre, n = 45) kindly donated from Professor Herbert Herzog (Garvan Institute, Sydney, [1, 2], Slc17a6$^{tm2(cre)Lowl}$/J (vGlut2-Cre, n = 16) obtained from Zane Andrews (Monash University) originally from the Jackson Laboratory (Bar Harbor, ME, USA, stock # 016963) were bred in house at the Florey Institute of Neuroscience and Mental Health. iCART-Cre mice were crossed with B6.Cg-Gt(ROSA)26Sor$^{tm14(CAG-TdTomato)Hze}$/J (*Ai14*) (Bar Harbor, ME, USA, stock # 007914) to produce iCART-*Ai14* offspring (n = 14). Mice were acclimatised to a reverse light cycle (lights off 7.00-19.00 AEST) and single housing to allow accurate measurement of fluid and food consumption. All mice had *ad libitum* access to food (laboratory chow, Barastoc) and water except where detailed. All studies were performed in accordance with the Prevention of Cruelty to Animals Act (2004), under the guidelines of the National Health and Medical Research Council (NHMRC) Council Code of Practice for the Care and Use of Animals for Experimental Purposes in Australia (2013) and approved by The Florey Animal Ethics Committee.

*Genotyping*

iCART-Cre, vGlut2-Cre and iCART-Ai14 mice had mutants identified using polymerase chain reaction (PCR) procedures provided by the supplier ([1, 2]; Jackson Laboratories).

*Stereotaxic surgery*

For surgeries, mice were anesthetized with 4% isoflurane and placed in a stereotaxic frame (Stoelting) and maintained under 1-2% isoflurane for the duration of the surgery. The skull was exposed, and burr holes were made with a drill over the coordinates of the EW (M/L + 1.00, A/P -3.40, D/V -4.70). For viral surgeries a Nanoject III (Drummond Scientific, Broomall, PA, USA) was used to deliver 100nL (50 nL/min) of an AAV virus. For DREADD experiments iCART-cre mice received AAV2-hsyn-DIO-hM4Di-mcherry or control vector AAV2-hsyn-DIO-mcherry (Addgene #44362-AAV2, # 50459-AAV2). For EW Ghsr knockdown C57BL6J mice received AAV2-hsyn-Ghsr-ShRNA-mCherry or scramble control, AAV2-hsyn-scramble-ShRNA-mCherry, while targeted deletion was achieved using Cre dependent viruses (AAV2-hSyn-DIO-Ghsr-ShRNA-mCherry or AAV2-hSyn-DIO-scramble-ShRNA-mCherry) administered into iCART-Cre or vGlut2-Cre mice (Vector Biolabs, Malvern, PA, USA). To activate the CART promoter controlled by the Cre-recombinase gene, and therefore induce specific expression of viruses into EW$^{CART}$ cells, tamoxifen (80 mg/kg, 10 ml/kg, i.p.) (Sigma-Aldrich Pty Ltd, Sydney, NSW, Australia) dissolved in 10% absolute ethanol and 90% sunflower oil was administered to CART-Cre mice for 5 consecutive days following surgery.

To allow direct drug administration to the EW, mice underwent stereotaxic cannula implantation. Mice were anaesthetized and prepared as per viral injections. Next, two holes were drilled into the skull and screws (Mr Specs. Parkdale, Australia) were inserted to anchor the cannula to the skull. A small hole was drilled into the skull, through which 26 G stainless-steel guide cannulas cut 3.5 mm below the pedestal (PlasticsOne,

Roanoke, VA, USA) were implanted into the EW (M/L + 1.00, A/P -3.40, D/V -3.20 mm on a 15 deg angle [3]. Cannula were fixed to the skull using dental cement (Vertex-Dental) and a dummy, cut projecting 1 mm beyond the cannula tip was inserted (PlasticsOne). Mice received meloxicam (3 mg/kg, i.p.) for analgesia, the antibiotic, baytril (3 mg/kg, i.p.) and saline (10 ml/kg, s.c.) at the end of surgery and were monitored for 5 days following surgery.

*Drugs*

LEAP2 (37-76) (5 μg, Phoenix Pharmaceuticals #075-58) or JMV2959 (10 μg, MedChemExpress, #HY-U00433A) [4] diluted in 10% DMSO in 0.9% sterile saline. Ghrelin (BOC Sciences, 1 mg/kg i.p.) was diluted in 0.9% sterile saline. CNO (10 mg/kg) was diluted in 0.05% DMSO in 0.9% sterile saline. All drugs were administered at 10 ml/kg.

*Drug microinfusion*

Intra-EW infusions of Vehicle, LEAP2 or JMV2959 were made using 40 cm polyethylene connectors (PlasticsOne) attached to 1 μL syringes (SGE Analytical Science, Ringwood). Mice were gently held while dummies were removed, and injectors inserted into the cannula. Compounds were infused (0.25 μL/min, 0.5 uL total) by an automated syringe pump (Harvard Apparatus, Holliston, USA). The injectors were left in place for ~2 min after infusion. Mice were habituated to the equipment prior to testing and each mouse received both drugs and vehicle treatment in a randomised and counterbalanced manner. Following behavioural testing to assess cannula positioning, methylene blue (0.5 μL) was infused and animals were anaesthetized with pentobarbitone (100 mg/kg, i.p. Virbac, Milperra, Australia) before decapitation. Brains were collected, frozen over super-cooled isopentane, sectioned (40 μm) on a cryostat (Leica, Leica Microsystems) and mounted onto SuperFrost Plus slides (Menzel-Glaser). After drying, slides were counterstained with Neutral Red solution (Sigma-Aldrich) for 2 min and cleared through a series of ethanol (50%, 70%, 90%, 100%) and X-3B (Olichem Pty Ltd.) washes before cover-slipping with SafteyMount mounting media (Sigma-Aldrich). An investigator blinded to treatment groups and behavioural outcomes performed injection site validations.

**Behaviour**

Binge drinking

*Training*

We adapted the SHAC (scheduled high access consumption) and DID (drinking in the dark) procedures. Mice had voluntary access to 10% (v/v) ethanol or 5% (w/v) sucrose diluted in tap water, 3 h after the dark cycle began, 3 days per week (Monday, Wednesday, Friday) for 2 h [5]. For DREADD experiments, mice received surgery prior to training and underwent 10 sessions of training before testing. For shRNA and cannulation studies, mice were trained for 6 sessions prior to surgery and given at least 4 sessions post-surgery before testing. To assess ghrelin induced drinking, mice were given access to alcohol in their light phase (3 h after the light cycle began, 3 days per week (Monday, Wednesday, Friday) for 2 h).

Given that ghrelin levels peak in the dark phase, to assess ghrelin-induced drinking, access to alcohol was given during the light phase, when ghrelin levels are low [6]. Mice had access to alcohol 3 h into the light cycle, for 2 h, 3 times weekly during training.

Mice were tested in an extended 4-hour drinking session with consumption measured hourly. Mice were given another session to ensure consumption returned to baseline, before testing with the counterbalanced treatment. For DREADD studies CNO was administered 1 h prior to testing. For ghrelin-induced drinking, ghrelin was administered 20 minutes prior to test onset, and food was removed for the duration of the test. Mice were weighed before binge drinking sessions and ethanol and sucrose consumption in grams per kilogram (g/kg) was calculated using the volume consumed in millilitres (ml) and body weight of each mouse in grams (g).

*Other behaviours in hM4Di mice*

During DREADD inhibition test sessions, food consumption was assessed by weighing standard chow at each timepoint after CNO administration. In a separate test, saccharin preference was assessed in a 4 h two bottle choice test where mice had access to either saccharin (0.01%) or water. Preference and intake were measured. To assess anxiety-like behaviours a light-dark box was employed [5]. Finally given the implication of the EW in body temperature regulation [7, 8], we assessed rectal temperature pre- and post-CNO administration.

*qPCR*

Mice were euthanised via cervical dislocation following behavioural testing. Brains were immediately removed and quickly frozen over liquid nitrogen until sectioning. The EW and VTA of each animal were dissected using needles with inner diameters 1.0-1.5 mm. RNA extraction and analysis were performed as previously described [9, 10]. Briefly, RNA from brain tissue was extracted using the QIAGEN Qiazol + RNeasy Plus Micro Kit according to the manufacturer's protocol. Total RNA (300 ng) was then reverse-transcribed into cDNA using SuperScriptII qRT-PCR kit (Invitrogen, USA). SYBR Powerup PCR kit (Qiagen) was used in 10 µL reactions containing 4 µL cDNA, and 1 µL 20 nmol/µL primer mixture. qPCR was performed with ViiA7(Applied Biosystems, USA), followed by melt-curve analysis. The qRT-PCR program was set to 2 min at 50 °C, 2 min at 95 °C, and 15 s at 95 °C and 1 min at 60 °C (40 cycles). Reactions were performed in 3 technical replicates along with a no template and water control. Two reference genes (*Actin* and *Hprt*) were used. Primers (Table S1) for were designed using Primer3 v. 0.4.0 software (http://frodo.wi.mit.edu/cgi-bin/primer3/primer3_www.cgi; Whitehead Institute for Biomedical Research, USA) based on coding DNA sequences acquired from GenBank (National Center for Biotechnology Information (NCBI)). Primer specificity was verified (BLAST Interface, NCBI)).

*Immunohistochemistry*

For viral injection sites: Mice were anaesthetised (pentobarbitone, 80 mg/kg, i.p.) and transcardially perfused with 15 mL PBS (0.1 M, pH 7.4) followed by 30 mL 4% PFA in PBS, decapitated, brains removed and post-fixed (1 h) in 10 mL perfusion solution. Brains were then incubated in 30% sucrose in PBS (10 mL) at 4°C overnight, before being frozen over liquid nitrogen and stored at −80$^{o}$C until sectioning. Coronal sections (40 μm) were cut on a cryostat (Leica Biosystems) at −18$^{o}$C and stored in PBS-azide in a 1 in 4 series. Fluorescent immunohistochemistry was performed to examine the co-localization of CART and viral reporter expression [11, 12]. Briefly, sections were blocked in 10% normal donkey serum (NDS) and 0.5% tritonX-100 (TX-100) in PBS, for 1 h at RT. Sections were then incubated in a primary antibody solution containing rabbit anti-CART (1:2000; Phoenix Pharmaceuticals, #H-003-62) and Chicken anti-RFP (Rockland #600-901-379) with 2% NDS in 0.1M PBS containing 0.1% TX-100 overnight at RT. Sections were washed 3 x 5 min in PBS and incubated for 2 h at room temperature in donkey anti-rabbit AlexaFluor488 (1:400; Life Technologies, #A-21206) and donkey anti-Chicken AlexaFluor594 (Jackson Immunology #703-585-155). Sections were finally washed (3 x 5 min) in PBS, mounted on microscope slides and coverslipped with fluorescence mounting medium (DAKO, Australia).

For electrophysiology slices tissue was washed in PBS for 3 x 5 minutes before antigen retrieval was undertaken. Tissue was incubated in wells of buffer (10mM Sodium Citrate, 0.05% Tween20, pH 6.0) in an oven at 90°C oven for 3 hours, then left to cool to room temp for 30 minutes. Tissue was washed in PBS for 3 x 5 minutes before blocked for 1 hour at room temp in 10% NDS + 0.3% TX-100 in PBS. Anti-CART h/m/r antibody (#AF163) at 1:1000 in 3% NDS + 0.3% TX-100 in PBS was next incubated with the tissue for 48 hours at 4°C with gentle agitation. Tissue was washed in PBS for 3 x 5 minutes before secondary antibody incubation. Streptavidin HRP 488 (#S11223, 1:500) and donkey anti-goat AlexaFluor594 (#A11058, Lot 1975275, 1:400), in 10% NDS + 0.3% TX-100 + PBS were incubated with tissue for 2 hat RT. Finally, tissue was washed in PBS for 3 x 5 minutes before mounting on slides with DAKO mounting media and imaging.

*In Situ Hybridisation*

To examine *Ghsr* expression across CART and vGlut2 cells in the EW, mice underwent cervical dislocation, brains were rapidly extracted and frozen for 20 secs on dry ice-cooled 2-methylbutane (Bacto Laboratories, NSW, Australia). Brains were stored at -80°C until use. Coronal sections of the EW (16 μm) were cut in a 1/6 series using a cryostat (Leica Biosystems, NSW, Australia) and mounted directly onto Super Frost Plus slides (Fisher Scientific, NH, USA). Slides were stored at -80°C until use. The RNAscope Multiplex Fluorescent Reagent Kit (Advanced Cell Diagnostics) was used to detect *Ghsr*, and *Slc17a6* (vGlut2) *and Cartpt* in the EW [13]. Slides were fixed in 4% w/v paraformaldehyde for 15 mins at 4°C, before rinsing in 0.1 M PBS (pH 7.4) followed by dehydration in increasing concentrations of ethanol (50%, 70%, 100% and 100% v/v) for 5 mins each at RT. Slides were then air-dried for 10min at RT and a hydrophobic barrier was drawn around the brain sections. Sections were then protease treated (pre-treatment 4) at RT for 5 min. Next, sections were rinsed in distilled water followed by probe application. The target probes used were, *Ghsr* (#426148), *Cartpt* (#432008-C2), *Slc17a6* (#319171-C3). Slides were first incubated with probes at 40°C for 1 hr. Following this, slides were incubated with preamplifier and amplifier probes (AMP1, 40°C for 30min; AMP2, 40°C for 15min; AMP3, 40°C for 30 min), followed by incubation in fluorescently labelled probes to select a specific combination of reporters associated with each channel (AMP4 Alt A to detect *Ghsr* in Alexa-488,

*Slc17a6* in Atto 550 channel and *Cartpt* in Atto 647). Finally, sections were incubated for 20 s with DAPI and coverslipped with DAKO Fluorescent Mounting Medium (North Sydney, NSW, Australia).

*Microscopy*

A LSM780 Zeiss Axio Imager 2 confocal laser scanning microscope (Carl Zeiss AG, Jena, Germany) with a 40 X objective was used to take all images of the EW (pixel size 0.10um). FISH analysis was quantified from at least 3 sections per mouse (~Bregma -3.5 mm; [3] using image J (National Institutes of Health; RRID:SCR_003070). For a cell to be deemed positive for the expression of a given gene, five or more mRNA puncta needed to be present [9, 12]. A LSM900 Zeiss Axio Imager 2 confocal laser scanning microscope was used to take images of sections used for electrophysiology and imaged at 10 x magnification.

*Electrophysiology*

Mice were deeply anesthetized using isoflurane, decapitated and brains were rapidly removed and mounted in a slice chamber containing chilled cutting solution (125 mM choline chloride, 20 mM D-Glucose, 0.4 mM $CaCl_2 \cdot 2H_2O$, 6 mM $MgCl_2 \cdot 6H_2O$, 2.5 mM KCl, 1.25 mM $NaH_2PO_4$ and 26 mM $NaHCO_3$). Slices were cut using a vibratome (Leica VT1200 S), and incubated in a holding chamber with oxygenated artificial cerebrospinal fluid (125 mM NaCl, 10 mM D-Glucose, 2 mM $CaCl_2 \cdot 2H_2O$, 2 mM $MgCl_2 \cdot 6H_2O$, 2.5 mM KCl, 1.25 mM $NaH_2PO_4$, and 26 mM NaHCO3) initially at 32°C for 30 min and then room temperature for at least 30 min before being transferred to the recording chamber.. Whole-cell patch-clamp recordings were obtained at 30°C from visually identified fluorescent CART neurons in the EW. Borosilicate glass electrodes (4–7 MΩ) were filled with an internal solution comprising 125 mM K-gluconate, 5 mM KCl, 2 mM MgCl2·6H2O, 10 mM HEPES, 4 mM ATP-Mg, 0.3 mM GTP-Na, 10 mM phosphocreatine, 0.1 mM EGTA, and 0.2% biocytin, with a pH of 7.2 and osmolarity of 290 mOsm. Baseline data were captured during aCSF bath application, and this was followed by continuous perfusion of aCSF containing Ghrelin (1 µM), with cell recordings commencing after 5 mins. Current- and voltage-clamp recordings were performed using an Axon Multiclamp 700B amplifier (Molecular Devices), Digidata 1440 digitizer (Molecular Devices), and pCLAMP version 10 software. Data were sampled at 50 kHz with a low-pass filter at 10 kHz.

All slice electrophysiology data were analysed using Clampfit version 10.7. Passive membrane properties were calculated from holding potentials of −70 mV and -50mV using −10 mV test pulses. All other slice electrophysiology data were collected in current clamp. Bridge balance and pipette capacitance neutralisation were manually applied throughout current-clamp experiments. Gap-free recordings (30s recording time) in the absence of holding current were conducted to measure resting potential and spontaneous firing properties such as firing frequency (Hz). Morphological characteristics were analysed from the first firing event that occurred during the 30 Is recording time. Threshold was defined as the voltage at which the rising membrane potential slope exceeded 20 mV/ms. Amplitude was defined as the AP peak relative to a normalized baseline averaged from 0 ms to 10 ms before threshold. Rise time was defined as the time from 10% to 90% of peak. Width was measured at 50% of peak height. Decay time was defined as the time from 100% to 50% of peak.

*Statistical Analysis*

The effects of Treatment or Genotype across Time were analysed using an analysis of variance (ANOVA), with repeated measures where appropriate. *Post-hoc* Bonferroni corrections were performed when appropriate statistical significance was achieved. For analysis of the effect of Treatment, Genotype or Sex only on behaviour, gene expression or electrophysiological properties, data were analysed using two-tailed Student's t-test. Pearson correlations were used to compare behaviour to gene expression data.

**References:**


1. Lau, J., et al., *CART neurons in the arcuate nucleus and lateral hypothalamic area exert differential controls on energy homeostasis.* Molecular metabolism, 2018. **7**: p. 102-118.
2. Lau, J., Y.-C. Shi, and H. Herzog, *Temperature dependence of the control of energy homeostasis requires CART signaling.* Neuropeptides, 2016. **59**: p. 97-109.
3. Paxinos, G. and K.B. Franklin, *Paxinos and Franklin's the mouse brain in stereotaxic coordinates*. 2019: Academic press.
4. Prieto-Garcia, L., et al., *Ghrelin and GHS-R1A signaling within the ventral and laterodorsal tegmental area regulate sexual behavior in sexually naïve male mice.* Psychoneuroendocrinology, 2015. **62**: p. 392-402.
5. Maddern, X.J., et al., *Cocaine and amphetamine regulated transcript (CART) mediates sex differences in binge drinking through central taste circuits.* Neuropsychopharmacology, 2024. **49**(3): p. 541-550.
6. Bodosi, B., et al., *Rhythms of ghrelin, leptin, and sleep in rats: effects of the normal diurnal cycle, restricted feeding, and sleep deprivation.* American Journal of Physiology-Regulatory, Integrative and Comparative Physiology, 2004. **287**(5): p. R1071-R1079.
7. Zuniga, A., A.E. Ryabinin, and C.L. Cunningham, *Effects of pharmacological inhibition of the centrally-projecting Edinger-Westphal nucleus on ethanol-induced conditioned place preference and body temperature.* Alcohol, 2020. **87**: p. 121-131.
8. Zuniga, A., et al., *Vesicular glutamate transporter 2-containing neurons of the centrally-projecting Edinger-Westphal nucleus regulate alcohol drinking and body temperature.* Neuropharmacology, 2021. **200**: p. 108795.
9. Walker, L.C., et al., *Acetylcholine muscarinic M4 receptors as a therapeutic target for alcohol use disorder: converging evidence from humans and rodents.* Biological Psychiatry, 2020. **88**(12): p. 898-909.
10. Walker, L.C., et al., *Muscarinic M4 and M5 receptors in the ventral subiculum differentially modulate alcohol seeking versus consumption in male alcohol-preferring rats.* British journal of pharmacology, 2021. **178**(18): p. 3730-3746.
11. Walker, L.C., et al., *Nucleus incertus corticotrophin-releasing factor 1 receptor signalling regulates alcohol seeking in rats.* Addiction Biology, 2017. **22**(6): p. 1641-1654.
12. Walker, L.C., et al., *Central amygdala relaxin-3/relaxin family peptide receptor 3 signalling modulates alcohol seeking in rats.* British journal of pharmacology, 2017. **174**(19): p. 3359-3369.
13. Wang, F., et al., *RNAscope: a novel in situ RNA analysis platform for formalin-fixed, paraffin-embedded tissues.* The Journal of molecular diagnostics, 2012. **14**(1): p. 22-29.
