## Supplemental Data for "Edinger-Westphal ghrelin receptor signalling regulates binge alcohol consumption in a sex specific manner"

Pearl et al.,

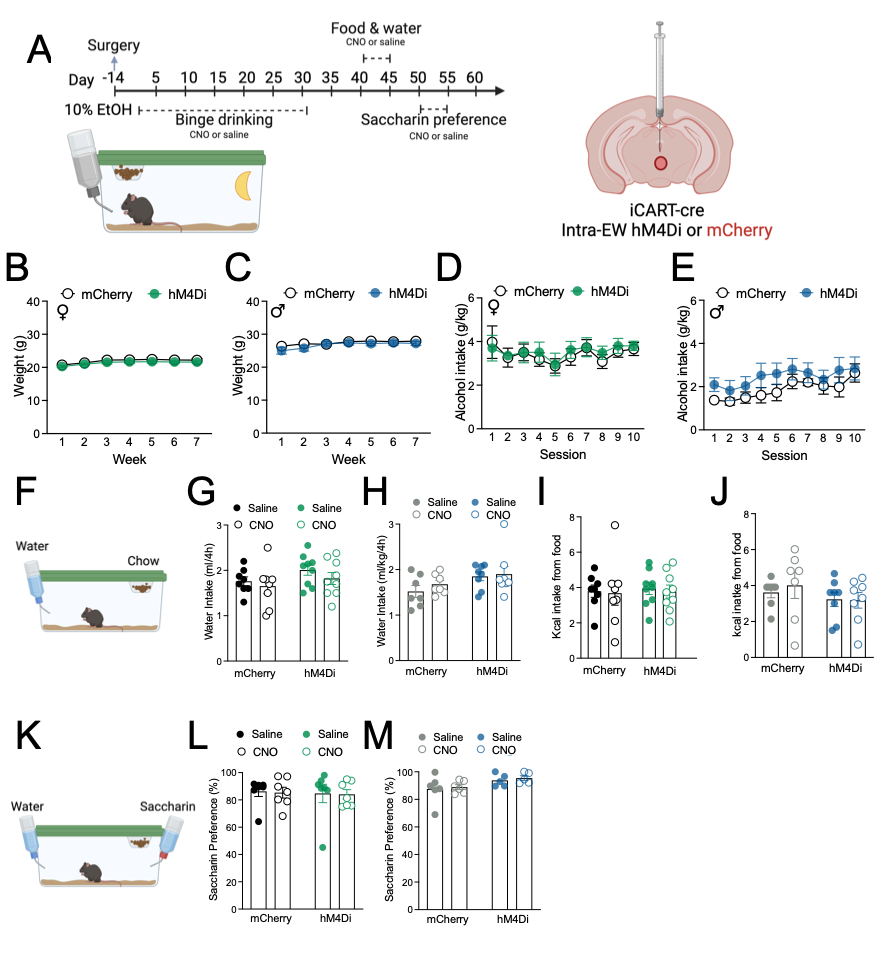
**SUPPLEMENTAL FIGURES**

**Supplementary Figure 1, related to Figure 1: Chemogenetic inhibition of EW^CART^ cells does not alter weight, water, food or saccharin intake. (A)** Schematic of experimental outline. RM two-way ANOVA showed no difference in weight of **(B)** female (virus F (1, 15) = 0.8869, p=0.3612; or interaction F (6, 90) = 0.2192, p=0.9697; but a main effect of time F (6, 90) = 24.09, p<0.0001), or **(C)** male mice (virus F (1, 13) = 0.4360, p=0.5206; or Interaction F (6, 78) = 1.852, p=0.0998; but a main effect of time (F (6, 78) = 16.80, P<0.0001) between HM4Di and control virus treated mice. Further during training no difference in alcohol consumption was observed in **(D)** female (RM two-way ANOVA, virus F (1, 15) = 0.1544, p=0.6999; interaction F (9, 135) = 0.1727, p=0.9965; or time F (9, 135) = 1.303, p=0.2407) or **(E)** male mice (RM two-way ANOVA, virus F (1, 13) = 1.217, P=0.2899; or interaction F (9, 117) = 0.6507, p=0.7515; but a main effect of time F (9, 117) = 6.612, p<0.0001) treated with HM4Di or control virus treated mice . **(F)** Schematic of food intake. No difference in water intake was observed in **(G)** female (RM two-way ANOVA, CNO F (1, 15) = 1.814, p=0.1980; virus F (1, 15) = 1.979, p=0.1799; or interaction F (1, 15) = 0.1287, p=0.7248) or **(H)** male mice (RM two-way ANOVA, CNO F (1, 13) = 0.9864, p=0.3387; virus F (1, 13) = 3.440, p=0.0865; or interaction F (1, 13) = 0.2832, p=0.6036) after administration of either CNO or vehicle. Further no difference in Kcal food intake was observed in **(I)** female (RM two-way ANOVA, CNO F (1, 15) = 0.1390, p=0.7145; virus F (1, 15) = 0.05401, p=0.8194; or interaction F (1, 15) = 0.004722, p=0.9461) or **(J)** male mice (RM two-way ANOVA, CNO F (1, 13) = 0.2711, p=0.6114; virus F (1, 13) = 1.017, p=0.3317; or interaction F (1, 13) = 0.4660, p=0.5068) after administration of CNO compared to vehicle. **(K)** Schematic of saccharin preference. No difference in 0.1% saccharin preference was observed in **(L)** female (RM two-way ANOVA, CNO F (1, 12) = 0.03693, p=0.8508; virus F (1, 12) = 0.06909, p=0.7971; or interaction F (1, 12) = 0.001049, p=0.9747) mice. **(M)** Male mice with hM4Di DREADD showed greater intake than mCherry mice (main effect virus F (1, 9) = 5.219, p=0.0482), however, there was no effect of CNO (F (1, 9) = 0.3088, p=0.5920; or interaction F (1, 9) = 0.003404, p=0.9548). Data expressed as mean ± SEM. n = 7-9 females/group and 5-8 male/group.

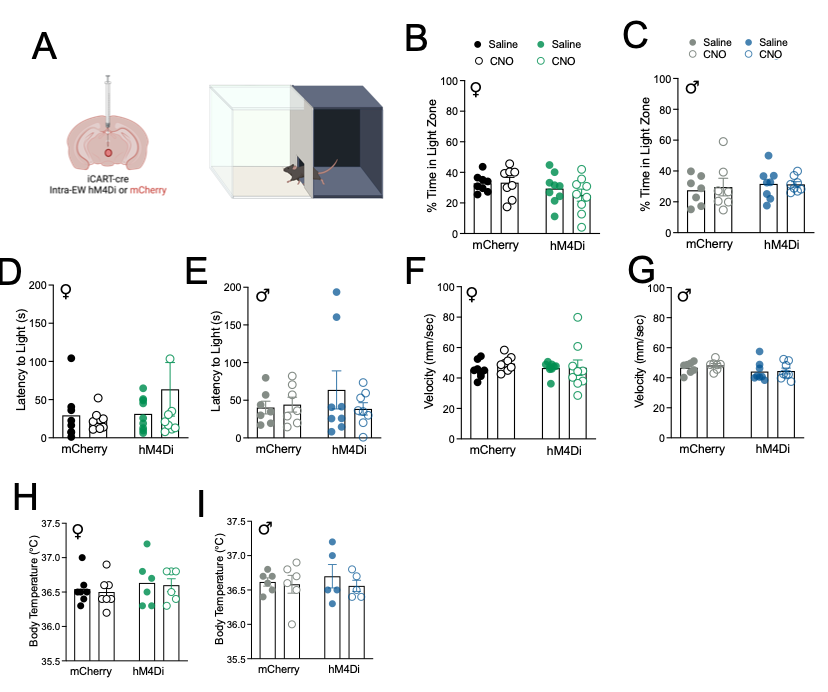
**Supplementary Figure 2, related to Figure 1: Chemogenetic inhibition of EW^CART^ cells does not alter anxiety-like behaviour or body temperature. (A)** Schematic of experiment outline of light-dark box. Two-way repeated measures ANOVA showed chemogenetic inhibition of EW^CART^ cells did not alter % time spent in the light zone of **(B)** female (CNO F (1, 15) = 0.4651, p=0.5056; virus F (1, 15) = 2.394, p=0.1426; or interaction F (1, 15) = 0.6733, p=0.4248) or **(C)** male control of DREADD treated mice (CNO F (1, 13) = 0.1344, p=0.7198; virus F (1, 13) = 0.3790, p=0.5488; or interaction F (1, 13) = 0.2504, p P=0.6252); latency to light in **(D)** female (CNO F (1, 15) = 0.4883, p=0.4954; virus F (1, 15) = 0.9889, p=0.3358; or interaction F (1, 15) = 1.025, p=0.3273) or **(E)** male control of DREADD treated mice (CNO F (1, 13) = 0.4987, p=0.4926; virus F (1, 13) = 0.3208, p=0.5807; or interaction F (1, 13) = 0.9266, p=0.3533). Further RM two-way ANOVA showed no effect of CNO on velocity in **(F)** female (CNO F (1, 15) = 0.4294, p=0.5222; virus F (1, 15) = 0.08511, p=0.7745; or interaction F (1, 15) = 0.3438, p=0.5664) or **(G)** male control or DREADD treated mice (CNO F (1, 13) = 0.2324, p=0.6378; virus F (1, 13) = 3.320, p=0.0915; or interaction F (1, 13) = 0.09966, p=0.7572. CNO has no effect on body temperature in **(H)** female (RM two-way ANOVA, CNO F (1, 11) = 0.4206, p=0.5299; virus F (1, 11) = 0.5349, p =0.4798; or interaction F (1, 11) = 0.006572, p=0.9368) or **(I)** male control or DREADD treated mice (RM two-way ANOVA, CNO F (1, 9) = 1.819, p=0.2103; virus F (1, 9) = 0.03924, p=0.8474; or interaction F (1, 9) = 0.6890, p=0.4280). Data expressed as mean ± SEM. n = 6-9/group.

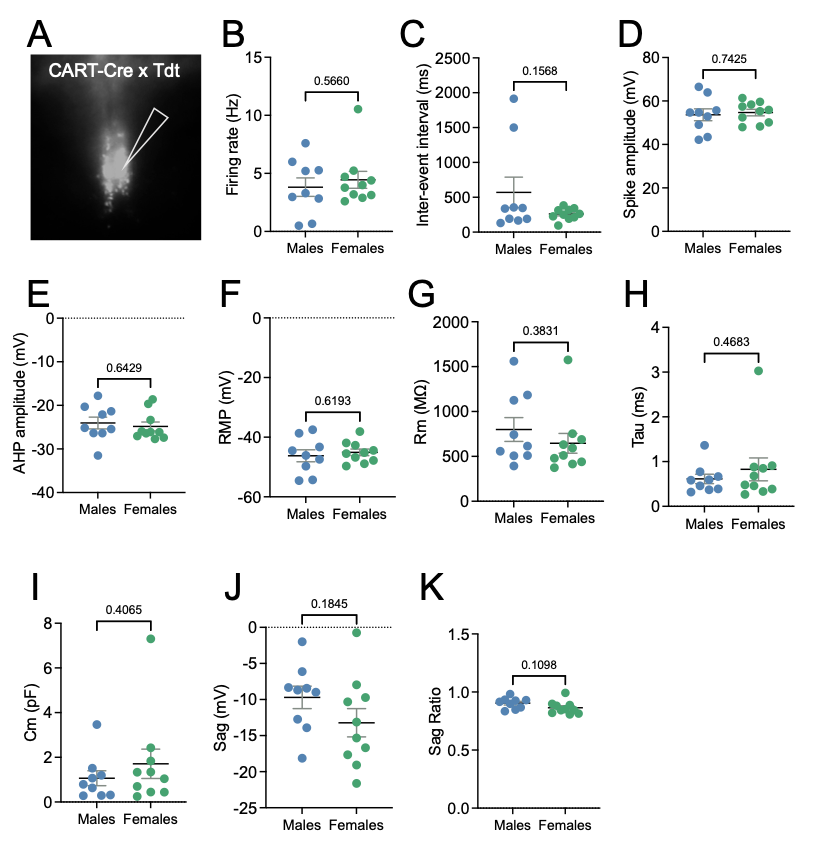
**Supplementary Figure 3, related to Figure 3: No sex difference in basal electrophysiological properties of EW^CART^ cells. (A)** overview of Tdtomato expression in the EW where patch clamp recording were collected. Unpaired t-test showed no sex differences were in **(B)** Firing rate (t=0.5854, df=17, p=0.5660), **(C)** inter-event interval (t=1.482, df=17, p=0.1568), **(D)** spike amplitude (t=0.3339, df=17, p=0.7425), **(E)** AHP amplitude (t=0.4720, df=17, p=0.6429), **(F)** resting membrane potential (RMP, t=0.5060, df=17, p=0.6193), **(G)** membrane resistance (Rm, t=0.8953, df=17, p=0.3831)**, (H)** tau (t=0.7418, df=17, p=0.4683), **(I)** membrane time constant (Cm, t=0.8511, df=17, p=0.4065), **(J)** sag (t=1.383, df=17, p=0.1845), or **(K)** sag ratio (t=1.687, df=17, p=0.1098) when directly comparing male and female mice. Data expressed as mean ± SEM. n = 7-10 cells from 5-7 mice/sex.

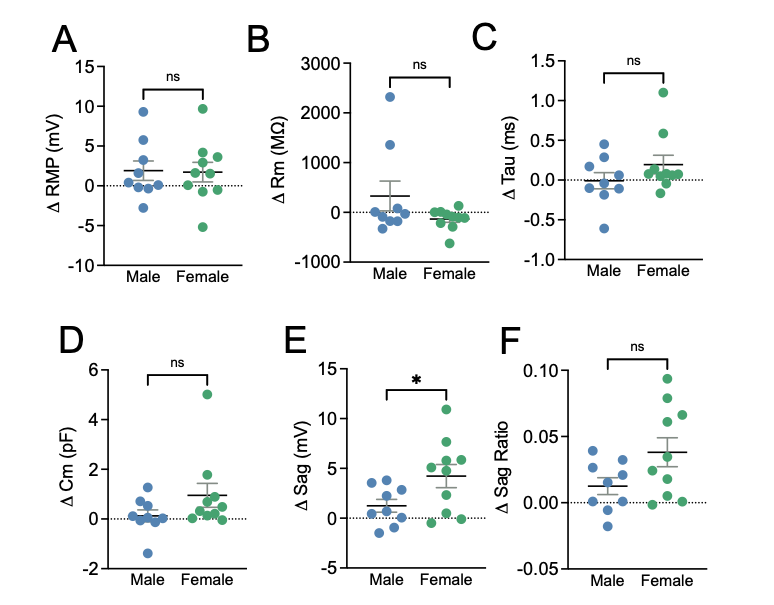
**Supplementary Figure 4, related to Figure 3: Sex differences in firing properties of EW^CART^ cells following ghrelin application.** Unpaired t-test showed no sex differences in ghrelin induced changes in **(A)** resting membrane potential (RMP, t=0.1051, df=17, p=0.9175), **(B)** membrane resistance (Rm, t=1.589, df=17,p= 0.1305), **(C)** tau (t=1.297, df=17, p=0.2118), **(D)** membrane time constant (Cm, t=1.479, df=17, p=0.1576), a significant difference in **(E)** sag (t=2.179, df=17, p=0.0437), and trend toward difference in **(F)** sag ratio (t=1.967, df=17, p= 0.0658) following bath application of ghrelin. Data expressed as mean ± SEM. n = 7-10 cells from 5-7 mice/sex.

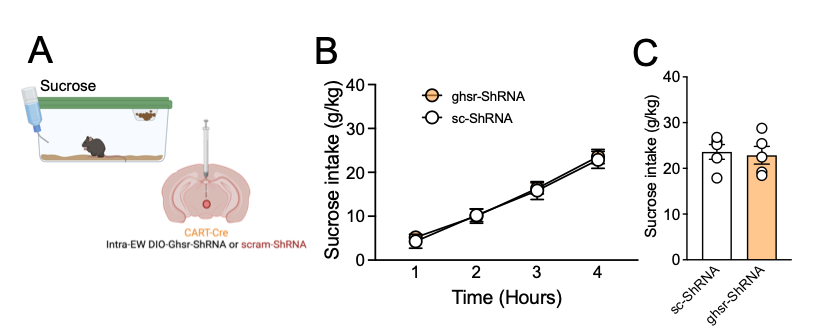
**Supplementary Figure 5, related to Figure 4: GHSR knockdown from EW^CART^ cells does not alter sucrose intake. (A)** Schematic of experiment outline of sucrose intake. No difference in **(B)** cumulative sucrose consumption over time (RM two-way ANOVA, virus F (1, 8) = 0.04907, p=0.8302; or interaction F (3, 24) = 0.5929, p=0.6257; but main effect of time F (3, 24) = 688.3, p<0.0001), or **(C)** total sucrose consumption (unpaired t-test, t=0.2934, df=8, p=0.7767) was observed in female mice with GHSR injected within the EW.

**SUPPLEMENTAL TABLES**

**Table S1: Key reagents**

| **Viruses** | **Supplier** | **Cat #** |
| --- | --- | --- |
| AAV2-hsyn-DIO-hM4Di-mcherry | Addgene | 44362-AAV2 |
| AAV2-hsyn-DIO-mcherry | Addgene | 50459-AAV2 |
| AAV2-hsyn-Ghsr-ShRNA-mCherry | Vector Biolabs | This paper |
| AAV2-hsyn-scramble-ShRNA-mCherry | Vector Biolabs | This paper |
| AAV2-hSyn-DIO-Ghsr-ShRNA-mCherry | Vector Biolabs | This paper |
| AAV2-hSyn-DIO-scramble-ShRNA-mCherry | Vector Biolabs | This paper |
| **Drugs** | **Supplier** | **Cat #** |
| LEAP2 (37-76) | Phoenix pharmaceuticals | #075-58 |
| JMV2959 | MedChemExpress, | #HY-U00433A |
| Ghrelin (i.p) | BOC Sciences | B0084-103854 |
| Ghrelin (bath) | BOC Sciences | B0084-103854 |
| **Antibodies** | **Supplier** | **Cat #** |
| rabbit anti-CART | Phoenix Pharmaceuticals | H-003-62 |
| Chicken anti-RFP | Rockland | 600-901-379 |
| Donkey anti-rabbit AlexaFluor488 | Life Technologies | A-21206 |
| Donkey anti-Chicken 594 | Jackson Immunology |  |
| **RNAscope** | **Supplier** | **Cat #** |
| RNAscope Multiplex Fluorescent Kit (V1) | Advanced Cell Diagnostics | 320850 |
| *Ghsr* | Advanced Cell Diagnostics | 426148 |
| *Cartpt* | Advanced Cell Diagnostics | 432008-C2 |
| *Slc17a6 (vGlut2)* | Advanced Cell Diagnostics | 319171-C3 |

**Table S2: Primer Sequences**

|  | **Forward** | **Reverse** |
| --- | --- | --- |
| ***Actin*** | GAACCCTAAGGCCAACCGTG | GGTACGACCAGAGGCATACA |
| ***Hprt*** | GCAGTACAGCCCCAAAATGG | GGTCCTTTTCACCAGCAAGCT |
| ***Ghsr*** | CTCAGGGACCAGAACCACAAAC | ACAAAGGACACCAGGTTGCAG |
| ***Esr1*** | CCTTGTCTCTTCCCTGATGTCAA | GTTCATTGTGACTGCCCTTGATC |
| ***Esr2*** | CGGTCTGTCTGAATGTGGTCA | GAAGCTGTGTGTGTGTGTGTC |
| ***Parq5*** | GTTACCGACACCCACAGAGTT | CGTCCAGATGTTGAGGGTCTC |
| ***Parq8*** | GACGACTGCCATCCTAGAGC | CTGCTGCCCACTCATTGACA |

**Table S3: Statistical analysis**

| **Table 3.1: Figure 1** | | | **Main effect** | | | | | |
| --- | --- | --- | --- | --- | --- | --- | --- | --- |
|  |  | **Analysis** | **Time** | | **Treatment** | | **Interaction** | |
| **d** | female | RM 2-way ANOVA | F (6, 90) = 24.09 | P<0.0001 | F (1, 15) = 0.8869 | P=0.3612 | F (6, 90) = 0.2192 | P=0.9697 |
|  | male | RM 2-way ANOVA | F (6, 78) = 16.80 | P<0.0001 | F (1, 13) = 0.6345 | P=0.3612 | F (6, 78) = 2.268 | P=0.9697 |
| **e** | female | RM 2-way ANOVA | F (9, 135) = 1.303 | P=0.2407 | F (1, 15) = 0.1544 | P=0.6999 | F (9, 135) = 0.1727 | P=0.9965 |
|  | male | RM 2-way ANOVA | F (9, 117) = 6.612 | P<0.0001 | F (1, 13) = 1.217 | P=0.2899 | F (9, 117) = 0.6507 | P=0.7515 |
| **f** |  | RM 2-way ANOVA | F (3, 21) = 450.9 | P<0.0001 | F (1, 7) = 2.116 | P=0.1891 | F (3, 21) = 0.5561 | P=0.6498 |
| **g** |  | Paired t-test |  |  | t=0.9523, df=7 | P=0.3727 |  |  |
| **h** |  | RM 2-way ANOVA | F (3, 27) = 184.6 | P<0.0001 | F (1, 9) = 19.45 | P=0.0017 | F (3, 23) = 2.523 | P=0.0828 |
| **i** |  | Paired t-test |  |  | t=3.378, df=8 | P=0.0097 |  |  |
| **j** |  | RM 2-way ANOVA | F (3, 18) = 150.8 | P<0.0001 | F (1, 6) = 0.5880 | P=0.4723 | F (3, 18) = 0.9155 | P=0.4532 |
| **k** |  | Paired t-test |  |  | t=0.1980, df=6 | P=0.8496 |  |  |
| **l** |  | RM 2-way ANOVA | F (3, 21) = 95.49 | P<0.0001 | F (1, 7) = 0.003947 | P=0.9517 | F (3, 21) = 0.5557 | P=0.6500 |
| **m** |  | Paired t-test |  |  | t=0.3474, df=7 | P=0.7385 |  |  |
| **n** | ctrl | Paired t-test |  |  | t=1.774, df=7 | P=0.1193 |  |  |
|  | hM4Di | Paired t-test |  |  | t=5.606, df=8 | P=0.0005 |  |  |
| **o** | ctrl | Paired t-test |  |  | t=1.413, df=6 | P=0.2073 |  |  |
|  | hM4Di | Paired t-test |  |  | t=0.9933, df=7 | P=0.3537 |  |  |
| **p** | ctrl | Paired t-test |  |  | t=0.09690, df=7 | P=0.9255 |  |  |
|  | hM4Di | Paired t-test |  |  | t=1.298, df=8 | P=0.2303 |  |  |
| **q** | ctrl | Paired t-test |  |  | t=0.5915, df=6 | P=0.5758 |  |  |
|  | hM4Di | Paired t-test |  |  | t=0.01623, df=7 | P=0.9875 |  |  |

| **Table 3.2: Figure 2** | | | **Main effect** | | | | | |
| --- | --- | --- | --- | --- | --- | --- | --- | --- |
|  |  | **Analysis** | **Time** | | **Treatment** | | **Interaction** | |
| **d** |  | RM 2-way ANOVA | F (2, 42) = 94.15 | P<0.0001 | F (2, 21) = 5.364 | P=0.0131 | F (4, 42) = 1.457 | P=0.2324 |
| **e** |  | RM 1-way ANOVA |  |  | F (2, 21) = 3.512 | P=0.0483 |  |  |
| **f** |  | RM 2-way ANOVA | F (1.385, 9.693) = 51.95 | P<0.0001 | F (1.825, 12.77) = 0.02530 | P=0.9674 | F (2.419, 16.93) = 0.1348 | P=0.9076 |
| **g** |  | RM 1-way ANOVA |  |  | F (2, 21) = 0.03941 | P=0.9614 |  |  |
| **j** | EW | unpaired t-test |  |  | t=5.729, df=23 | P<0.0001 |  |  |
|  | VTA | unpaired t-test |  |  | t=0.7605, df=22 | P=0.455 |  |  |
| **k** |  | RM 2-way ANOVA | F (8, 112) = 2.934 | P=0.0052 | F (1, 14) = 2.837 | P=0.1143 | F (8, 112) = 3.329 | P=0.0019 |
| **l** |  | RM 2-way ANOVA | F (1.993, 21.92) = 189.7 | P<0.0001 | F (1, 11) = 5.328 | P=0.0414 | F (3, 33) = 1.558 | P=0.2181 |
| **m** |  | unpaired t-test |  |  | t=2.527, df=11 | P=0.0281 |  |  |
| **n** |  | linear regression |  |  | R2 = 0.4332 | P=0.0145 |  |  |
| **o** |  | RM 2-way ANOVA | F (4.223, 40.65) = 2.716 | P=0.0404 | F (1, 10) = 3.236 | P=0.1022 | F (8, 77) = 1.201 | P=0.3096 |
| **p** |  | RM 2-way ANOVA | F (2.635, 26.35) = 3.584 | P=0.0315 | F (1, 10) = 3.632 | P=0.0858 | F (4, 40) = 1.635 | P=0.1843 |
| **q** |  | unpaired t-test |  |  | t=0.2184, df=10 | P=0.8315 |  |  |
| **r** |  | linear regression |  |  | R2 = 0.0001 | P=0.953 |  |  |

| **Table 3.3: Figure 3** | | |  |  |
| --- | --- | --- | --- | --- |
|  |  | **Analysis** | **stats** | **p value** |
| **d** | *Ghsr* | unpaired t-test | t=1.950, df=7 | P=0.0922 |
|  | *Cartpt* | unpaired t-test | t=0.7822, df=7 | P=0.4598 |
|  | *vGlut2* | unpaired t-test | t=1.207, df=7 | P=0.2668 |
| **g** |  | unpaired t-test | t=2.232, df=9 | P=0.0525 |
| **h** |  | unpaired t-test | t=3.850, df=9 | P=0.0039 |
| **i** |  | unpaired t-test | t=2.628, df=9 | P=0.0274 |
| **j** |  | unpaired t-test | t=1.296, df=9 | P=0.2273 |
| **k** |  | unpaired t-test | t=0.1361, df=9 | P=0.8947 |
| **m** |  | paired t-test | t=6.452, df=9 | P<0.0001 |
| **m** |  | paired t-test | t=5.614, df=8 | P=0.0005 |
| **n** |  | unpaired t-test | t=2.178, df=17 | P=0.0438 |
| **n** |  | paired t-test | t=8.425, df=9 | P<0.0001 |
| **o** |  | paired t-test | t=8.512, df=8 | P<0.0001 |
| **p** |  | unpaired t-test | t=0.9445, df=17 | P=0.3581 |

| **Table 3.4: Figure 4** | | | **Main effect** | | | | | |
| --- | --- | --- | --- | --- | --- | --- | --- | --- |
|  |  | **Analysis** | **Time** | | **Treatment** | | **Interaction** | |
| **d** |  | RM 2-way ANOVA | F (2.993, 47.88) = 4.752 | P=0.0056 | F (1, 16) = 9.113 | P=0.0082 | F (5, 80) = 0.9294 | P=0.4665 |
| **e** |  | RM 2-way ANOVA | F (1.664, 26.63) = 225.6 | P<0.0001 | F (1, 16) = 10.99 | P=0.0044 | F (3, 48) = 4.471 | P=0.0076 |
| **f** |  | unpaired t-test |  |  | t=3.833, df=16 | P=0.0015 |  |  |
| **j** |  | RM 2-way ANOVA | F (3.164, 53.78) = 1.317 | P=0.2783 | F (1, 17) = 0.002752 | P=0.9588 | F (5, 85) = 0.1803 | P=0.9693 |
| **k** |  | RM 2-way ANOVA | F (1.643, 27.92) = 188.0 | P<0.0001 | F (1, 17) = 0.8315 | P=0.3746 | F (3, 51) = 1.597 | P=0.2015 |
| **l** |  | unpaired t-test |  |  | t=1.247, df=17 | P=0.2293 |  |  |

| **Table 3.5: Figure 5** | | | **Main effect** | | | | | |
| --- | --- | --- | --- | --- | --- | --- | --- | --- |
|  |  | **Analysis** | **Time** | | **Treatment** | | **Interaction** | |
| **d** |  | RM Mixed effects | F (11, 120) = 1.829 | P=0.0562 | F (1, 11) = 0.2343 | P=0.6379 | F (11, 120) = 0.2228 | P=0.9957 |
| **e** | EW | Paired t-test |  |  | t=0.5968, df=6 | P=0.5725 |  |  |
|  | Ctrl | Paired t-test |  |  | t=0.8929, df=4 | P=0.4224 |  |  |
| **f** |  | RM 2-way ANOVA | F (1.471, 8.825) = 42.41 | P<0.0001 | F (1.431, 8.583) = 11.37 | P=0.0058 | F (2.760, 16.56) = 5.695 | P=0.0082 |
| **g** |  | RM 1-way ANOVA |  |  | F (1.625, 9.749) = 11.60 | P=0.0036 |  |  |
| **h** |  | RM 2-way ANOVA | F (1.270, 5.082) = 43.57 | P=0.0009 | F (1.748, 6.991) = 0.7170 | P=0.5029 | F (1.969, 7.876) = 1.674 | P=0.2476 |
| **i** |  | RM 1-way ANOVA |  |  | F (1.127, 4.510) = 1.004 | P=0.3797 |  |  |
